# Spatial distribution of Tree-related Microhabitats in a primeval mountain forest: from natural patterns to landscape planning and forest management recommendations

**DOI:** 10.1101/2024.04.25.591232

**Authors:** Fabian Przepióra, Paweł Lewandowski, Michał Ciach

## Abstract

Tree-related Microhabitats (TreMs) are essential for sustaining forest biodiversity. Although TreMs represent ephemeral resources that are spread across the landscape, their spatial distribution within temperate forests remains poorly understood. To address this knowledge gap, we conducted a study on 90 sample plots (0.05 ha each) located in a primeval mountain European beech *Fagus sylvatica*-dominated forest (Bieszczady Mountains, Carpathians). We explored the TreM profile with its link to habitat characteristics and described the spatial distribution of TreM indices. We identified 61 TreM types, with a mean richness of 19.7 ±4.9 SD TreM types per plot, a mean density of 740.7 ±292.5 SD TreM-bearing trees ha^-1^ and a mean TreM diversity of 1.2 ±0.1 SD. The diameter and living status of trees (living *vs* dead standing tree) were correlated with TreM richness on an individual tree. The stand structure, i.e. density and/or basal area of living and/or dead standing trees, and topographic conditions, i.e. slope exposure, were correlated with the TreM richness, density and diversity recorded on a study plot. We found no relationship between TreM richness, density and diversity and the presence of canopy gaps, which indicates that the influence of small-scale disturbances on the TreM profile is limited. However, our analysis revealed a clustered spatial pattern of TreM indices, with TreM-rich habitat patches (hot-spots) covering ∼20% of the forest. A moderate TreM richness, density and diversity dominated ∼60% of the forest, while TreM-poor habitat patches (cold-spots) covered ∼20%. Based on our findings, we advise the transfer of knowledge on the spatial distribution of TreMs from primeval to managed forests and advocate the ‘2:6:2’ triad rule: to allocate 20% of forests as strictly protected areas, to dedicate 60% to low-intensity forest management with the retention of large living trees and all dead standing trees, and to use the remaining 20% for intensive timber production. To ensure the continuance of the majority of TreM types, ≥ 55 living trees ha^-1^ > 60 cm in diameter should be retained. Such an approach will maintain a rich and diverse TreM assemblage across a broad spatial scale, which in turn will support biodiversity conservation and ecosystem restoration in secondary or managed forests.

**Highlights:**

- Primeval mountain beech forest hosts rich and abundant TreM assemblage
- Stand structure, slope exposure, but not canopy gaps mould TreM profiles
- Habitat patches with high/low TreM indices are spatially clustered
- Hot- and cold-spots each make up 20% of TreM richness, density and diversity
- 2:6:2 - the protected: managed forest ratio for maintaining the TreM assemblage

## 1. Introduction

The global loss, degradation and fragmentation of natural habitats is leading to a decline in biodiversity (Jaureguiberry et al., 2022). Among the most compromised terrestrial ecosystems are forests, which have been subject to human-induced transformations over the centuries (Williams, 2003). At present, more than 75% of primeval forests have been lost worldwide (Kormos et al., 2018) and their remnants, with much smaller populations of forest-related taxa, are scattered across landscapes (Haddad et al., 2015). Reduced habitat connectivity is an additional threat to the survival of habitat specialist species, especially those with limited movement capabilities (Fietz et al., 2014). Therefore, the persistence of primeval forest remnants and their connectivity efficiently sustained by wildlife corridors is critical for the conservation of forest-dwelling species if further biodiversity loss is to be prevented (Vandekerkhove et al., 2013; Dinerstein et al., 2020).

Habitats designated as cores of protected areas or as wildlife corridors should exhibit a high level of heterogeneity that will provide resources for a wide range of taxa (Bouget et al., 2014). A mixed tree species composition, an abundance of various types of coarse woody debris, a high level of structural complexity and a diverse microclimate are among the forest traits that increase habitat heterogeneity and support biodiversity (Ampoorter et al., 2019; Heidrich et al., 2020; Tinya et al., 2021). Another forest feature crucial for a high level of habitat heterogeneity and biodiversity is a rich, abundant and diverse assemblage of Tree-related Microhabitats (hereafter: TreMs), i.e. structures occurring on living or dead standing trees (hereafter: snags) that are key habitats for a great many birds, mammals, amphibians, reptiles and insects (Bütler et al., 2013). More than 60 distinct TreM types have been documented within temperate forests (Kraus et al., 2016), and each one offers a place for living, breeding, feeding or shelter for an array of species, some of which are rare or on the verge of extinction (Winter and Möller, 2008; Regnery et al., 2013; Paillet et al., 2018; Basile et al., 2020).

To sustain the spatial continuity of habitats for a diverse range of species, the composition of TreMs in secondary or managed forests should reflect that of natural ones (Vandekerkhove et al., 2013; O’Brien et al., 2021). For example, secondary cavity-nesting birds need a continuum of specific cavity types: some owl species require cavities with a large entrance, small birds need smaller entry holes (Newton, 1994; van der Hoek et al., 2017). Depending on the season and temperature, bats utilize cracks, scars or cavities on different parts of trees, opting for TreMs in higher canopy sites for rearing their young and moving to structures located in deeper, lower sections for hibernation during winter (Bat Tree Habitat Key, 2018). Furthermore, different insects living in water-filled tree-holes need a variety of structural, trophic and physicochemical features such as tree hole depth, or amounts of organic litter, water and oxygen (Schmidl et al., 2008). However, the numbers of specific TreM types required to create suitable habitats for the numerous assemblages of birds, bats or invertebrates are not clearly defined (Asbeck et al., 2021). One solution for addressing this knowledge gap is to determine the numbers of relevant TreMs in forest patches where a target species is present and to replicate these numbers in forests where the species is intended to occur (Bütler et al., 2004; Volis, 2019). However, little information is available about the reference levels of TreM numbers, diversity and composition, or on the spatial arrangements to be achieved in habitat patches under diverse conservation/management regimes.

The spatial distribution of TreMs has received surprisingly little attention from researchers, and no comprehensive description exists of the spatial patterns of TreM occurrence in forests (Martin et al., 2022). The numbers and diversity of TreMs depend on the distinctive tree species composition, the living status of trees, the stand structure and the disturbance history (Martin et al., 2021a; Przepióra and Ciach, 2023). As a result, the spatial distribution of TreMs in natural forests is potentially non-uniform. An understanding of the spatial variation of TreM richness, abundance and diversity, especially in old-growth forests, considered to be highly heterogeneous (Franklin and Van Pelt, 2004), would provide valuable guidance for forest managers or landscape planners seeking solutions that emulate natural processes within ecosystems (Kuuluvainen et al., 2021). For instance, a rich and diverse assemblage of TreMs can only develop in sufficiently large patches of forest with a specific set of structural traits and tree species composition (Larrieu et al., 2012; Larrieu et al., 2014). As some set-aside forest patches or clusters of habitat trees are more and more often designated for the protection of forest-dwelling species in core areas or in wildlife corridors (Baker et al., 2021; NaturScot, 2023), practical recommendations aiming to secure the near-natural profiles and spatial arrangements of TreMs are becoming indispensable.

The occurrence of TreMs may be influenced by local variations in topography and disturbance regimes. For instance, elevation and slope exposures influence insolation and interfere with local wind patterns (Johansson and Chen, 2003; Salzmann et al., 2007), leading to local disparities in precipitation and temperature, which further affect the occurrence of tree cavities (Remm and Lõhmus, 2011). Steep slopes exacerbate the risks of rock or snow avalanches, which can result in tree damage, increasing the prevalence of trees with bark loss or root buttress cavities (Homma, 1997; Šilhán, 2010; Pop et al., 2016; Larrieu et al., 2022). Small-scale disturbances caused by strong winds and heavy snowfall may kill a single tree or a group of them, resulting in the emergence of canopy gaps (Lewandowski et al., 2021), which may then become local concentrations of TreM-rich damaged, injured or dead standing trees (Hobi et al., 2015). Moreover, local variations in topography introduce diverse soil and hydrological conditions, and canopy gaps produced by small-scale disturbances increase the insolation (Hobi et al. 2015; Metzen et al. 2019). Rugged terrain and disturbances locally alter the stand structure and facilitate the appearance of new tree species in the species pool, enhancing heterogeneity on a larger scale (Bolstad et al. 2018; Ishii et al. 2004). Increased heterogeneity fosters the development of distinct forest patches, each characterized by unique stand structures and species compositions (Franklin and Van Pelt, 2004), potentially harbouring different TreM profiles (Przepióra and Ciach, 2023). Hence, understanding the interplay among the stand structure, topography, presence of canopy gaps and TreM occurrence is crucial for identifying habitat patches with the richest and most diverse TreM assemblages.

European beech *Fagus sylvatica* is one of the major deciduous tree species in Europe, playing a significant role in forest management and biodiversity conservation (Brunet et al., 2010). Although European beech-dominated forests cover as much as ca 8% of the continent’s total woodland area (Brunet et al., 2010; Eurostat, 2020), only a few stands of primeval character have survived to the present day that offer sufficiently large sites for studying the spatial distribution of TreM numbers and diversity (Sabatini et al., 2018). One region where such research is possible is the well-preserved European beech-dominated forest in the Bieszczady Mountains (Mts.) (northern Carpathians), a larger area of which has survived. The forests of the Bieszczady Mts. harbour numerous target species, including rare and endangered lichens, fungi, vertebrates and invertebrates (Witkowski et al. 2003; Kościelniak, 2009; Gierczyk et al. 2019), making them a suitable benchmark for habitat structure and quality in the restoration of TreM-dependent species in forests designated as core areas or wildlife corridors.

In this study, we aim (1) to describe the density and frequency of occurrence of particular TreMs and the richness, abundance and diversity of the TreM assemblage in a primeval montane European beech-dominated forest in the Carpathians; (2) to investigate the relationships between the forest stand structure, topographic conditions, the presence of canopy gaps and TreM assemblages at the scale of an individual tree and of the study plot; and (3) to analyse the spatial pattern of variation of TreM richness, density and diversity with the specific task of identifying patches of habitat with rich and poor TreM profiles. We expected to discover spatially diversified, rich and abundant TreM assemblages, shaped by the topography and stand characteristics of this primeval mountain forest, and which could act as a reference for secondary or managed forests.

## 2. Methods

### 2.1. Study area

The research was conducted in the Bieszczady Mts. (northern Carpathians) (Fig. 1a), which straddle a transboundary area, a significant proportion of which is protected by national parks. They include the Bieszczady National Park (BNP) in south-eastern Poland (Fig. 1b) (area: 29,202 ha; established in 1973), the Poloniny National Park in north-eastern Slovakia (29,805 ha; established in 1997), and the Uzhanian National Park in south-western Ukraine (39,159 ha; established in 1999). As a result, all three national parks constitute one of the largest complexes of protected areas in Europe, known as the Eastern Carpathians Transboundary Biosphere Reserve (Taggart-Hodge and Schoon, 2016).

**Fig. 1.**
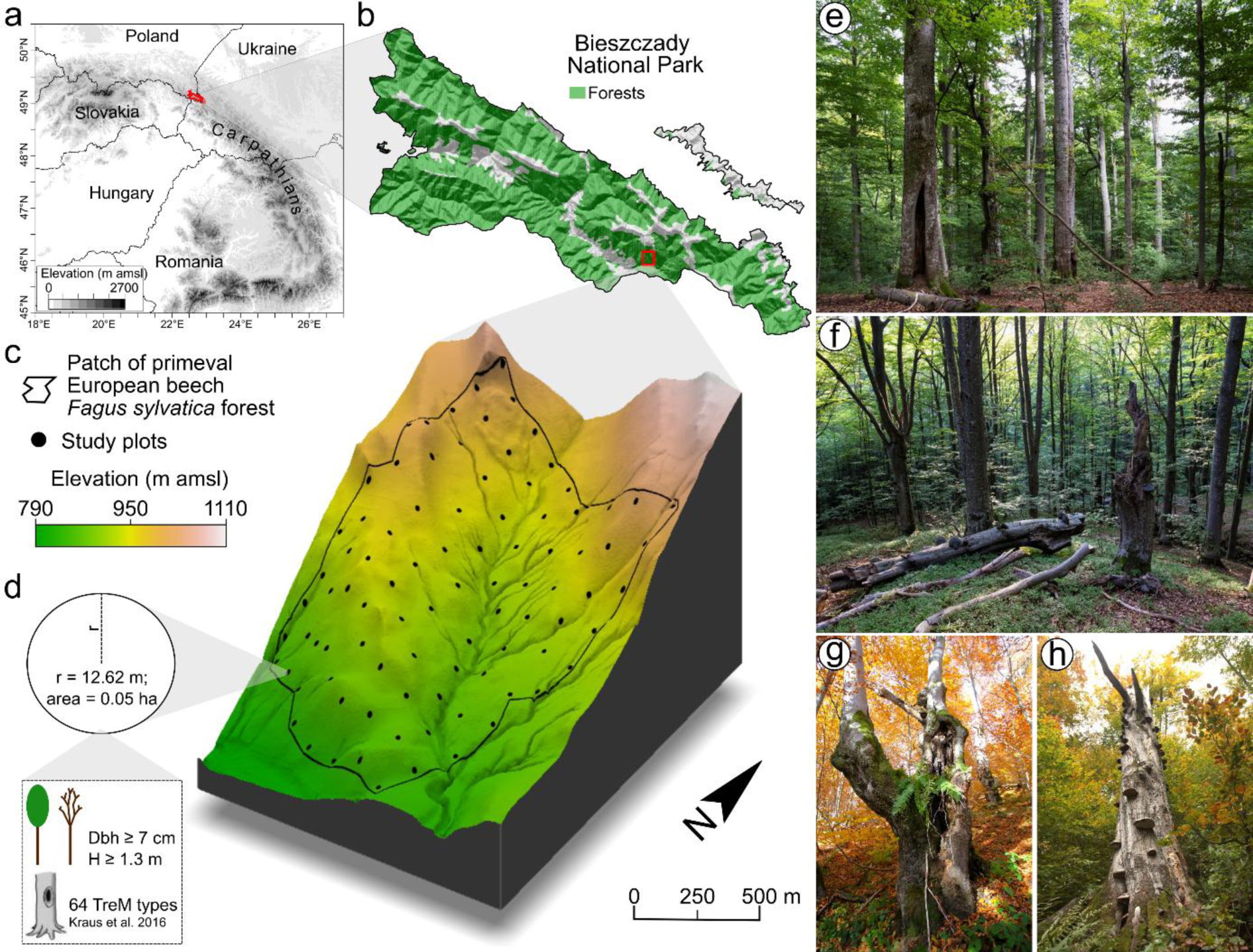
(a) Location of the study region – the Bieszczady National Park (BNP) covering the part of the Polish Bieszczady Mountains that straddles the transboundary area in the northern Carpathians, **(b)** location of the study area – the lower montane zone forests in the BNP, **(c)** location of 90 study plots within the boundaries of the primeval European beech *Fagus sylvatica*-dominated forest patch in relation to the topographic conditions and elevation (expressed in metres above mean sea level) of the area, **(d)** design of tree measurements and Tree-related Microhabitat (TreM) cataloguing performed on each study plot (in panel d, Dbh is the diameter at breast height (∼1.3 m) and H is the height of the tree; “TreM types” refers to the structures listed in Kraus et al., 2016), **(e)** primeval European beech-dominated forest in the study area with a wide variety of tree ages and a complex structure, **(f)** canopy gap with dead standing trees (snags) and downed logs initiating local tree regeneration, and examples of large **(g)** living trees and **(h)** snags hosting a rich diversity of TreMs (photos e-f by Paweł Lewandowski, photos g-h by Fabian Przepióra).

The mountains are characterized by a distinctive pattern of straight ridges aligned from north-west to south-east, separated by deep valleys, and with mean slope inclinations from 25 to 31° (Haczewski et al., 2007). The altitude of the highest peak in the Polish part of the Bieszczady Mts. is 1,346 m above mean sea level (amsl), while the bottoms of the lowest valleys lie at ca 400 m amsl. The elevation-related climate-vegetation zones in the Bieszczady Mts. are well defined, with the lower montane zone consisting mostly of European beech-dominated forests (75.6% of the forest area in BNP) that extend up to the upper tree line, which is situated at an elevation of ca 1,150 m amsl (Michalik and Szary, 2016).

Mean annual temperatures in the Bieszczady Mts. range from ca 6°C at elevations between 500-600 m amsl to ca 1°C at 1,200-1,300 m amsl (Nowosad, 1995). Mean monthly air temperatures range from 14°C in July to −5°C in January. The mean annual precipitation is 1,100-1,200 mm, and the area experiences an average of 135 days with continuous snow cover (Nowosad and Wereski, 2016). The prevailing winds in the study region are southerly and south-westerly (Wibig, 2021).

The study plots were located in the primeval European beech-dominated forest of the BNP, which is part of the UNESCO World Heritage Site “Ancient and Primeval Beech Forests of the Carpathians and Other Regions of Europe” that covers ca 100,000 ha of the best-preserved, strictly-protected European beech-dominated forests across 18 European countries (Kirchmeir and Kovarovics, 2023). The forests included in the UNESCO site cover ca 11% of the BNP’s area and have been preserved in the form of four patches several hundred hectares in area, situated in inaccessible, high and steep parts of the mountains (Kucharzyk, 2008). Historical cadastral maps from the 19^th^ century, the sparse population and the small number of logging roads, skid trails and timber transportation railways surrounding the study region, all indicate that the forest patch of ca 100 ha investigated in this study (Fig. 1c) has never been commercially exploited (Augustyn and Kucharzyk, 2008) and is one of the best-preserved parts of the Carpathian beech forests.

The remaining primeval forests in the BNP are dominated by European beech, with an admixture of sycamore *Acer pseudoplatanus*, silver fir *Abies alba* and Norway spruce *Picea abies* (Michalik and Szary, 2016); they also have a complex tree age- and size-structure (Kucharzyk, 2008) (Fig. 1e). Spatial and temporal changes of tree stands are driven primarily by canopy gap dynamics (Fig. 1f). Most canopy gaps come into existence after the death of an individual tree or a group of them, resulting in openings up to 0.1 ha in size (Lewandowski et al., 2021). The tree population includes large living trees (Fig. 1g) and large snags (Fig. 1h), and the average volume of living trees is ca 490 m^3^ ha^-1^, while the average volume of snags and downed logs is ca 55 m^3^ ha^-1^ (Kacprzyk et al., 2014). The forests in the BNP are inhabited by large mammals such as European bison *Bison bonasus*, red deer *Cervus elaphus* and brown bears *Ursus arctos*, which can damage individual trees through bark scratching or stripping (Broughton et al., 2022). The area hosts eight species of woodpeckers: great spotted woodpecker *Dendrocopos major*, white-backed woodpecker *Dendrocopos leucotos*, middle spotted woodpecker *Dendrocoptes medius*, lesser spotted woodpecker *Dryobates minor*, grey-headed woodpecker *Picus canus*, European green woodpecker *Picus viridis*, Eurasian three-toed woodpecker *Picoides tridactylus* and black woodpecker *Dryocopus martius* (Głowaciński, 2016; Lewandowski et al., 2021), all of which excavate cavities of various sizes. The area also hosts large saproxylic beetles, such as the rosalia longicorn *Rosalia alpina*, whose larval emergence holes are up to 1.5 cm in width (Ciach and Michalcewicz, 2013).

### 2.2. Field methods

The fieldwork was carried out in 90 circular study plots of radius 12.62 m (0.05 ha), which, within the boundaries of the selected patch of primeval European beech-dominated forest (Fig. 1c), were selected in a quasi-regular grid to maintain a minimum distance between the plots greater than double the stand height. The mean distance between the ultimately selected adjoining plots was 73.4 m ±12.9 SD (range 41-98 m). The fieldwork was carried out during the leafless period (March-May and October) in 2019. On each plot, every standing tree with a diameter at breast height, i.e. the diameter measured at a height of 1.3 m above the ground expressed in cm (hereafter, diameter), ≥ 7 cm was examined for the presence of TreMs (Fig. 1d). The species and living status of each tree (living tree *vs* snag) were identified, and the diameter and height of each one were measured.

TreMs were catalogued in accordance with the types described for temperate forests (Kraus et al., 2016), and cataloguing involved confirming the presence or absence of such a structure on a given tree (Przepióra and Ciach, 2022). TreMs were catalogued by two observers (the first two authors of this paper) who, prior to the fieldwork, were trained in carrying out a TreM inventory so as to minimize the observer effect (Paillet et al., 2015). Each tree was scrutinized for a minimum of 3 minutes, with the upper parts of the trunk and crown being examined through binoculars.

### 2.3. Remote-sensed data

The topographic characteristics, the occurrence of canopy gaps and the stand structure of the investigated forest patch (see Fig. 1c) were calculated using national open-source Light Detection and Ranging (LiDAR) data (obtained from airborne laser scanning with density of 4 points per m^2^; https://www.geoportal.gov.pl/en/data/lidar-measurements-lidar) and stored as a multi-layer raster with resolution 22.36 m (∼500 m^2^, similar to the size of the study plots). Before calculating the topographic characteristics, the raw LiDAR data were transformed into a Digital Elevation Model (DEM). The topographic characteristics were calculated for each cell in a raster layer and included elevation, slope inclination, Topographic Position Index (TPI), northness and eastness. Elevation, which is the height of a pixel in relation to mean sea level (expressed in m amsl), and slope inclination (expressed in degrees) were read indirectly from the DEM. The measure of terrain ruggedness – TPI – was calculated as the difference of elevation between a central pixel and the mean of its eight surrounding cells. Large negative values of TPI indicated a pixel as the bottom of a valley or gulley, large positive values indicated a pixel as the top of a ridge or hill, and values close to zero indicated a flat area (where the slope inclination is near zero) or a mountain slope (where the inclination is greater than zero) (Weiss, 2001). Northness was calculated as the cosine of the aspect with values between 1 – due north, and −1 – due south, with 0 being neither north nor south in aspect. Eastness was calculated as the sine of the aspect with values between 1 – due east, and −1 – due west, with 0 being neither east nor west in aspect.

Prior to calculating the canopy gap-related characteristics, the raw LiDAR data were transformed into a Canopy Height Model (CHM) with a resolution of 0.5 m. The canopy gaps were automatically detected based on local differences in the height of the CHM pixels measured by a moving search window (Silva et al., 2019). The minimum area of a detected gap was set at 0.03 ha, which corresponds to the mean area of canopy gaps recorded in the study area region (Lewandowski et al., 2021), and the maximum at 5 ha, which corresponds to the largest mid-forest open areas recorded close to the studied forest patch. The threshold difference in canopy height between contiguous areas was set at 10 m, which is the value adopted for research into canopy gaps (Silva et al., 2019). The detected canopy gaps were stored as a vector layer, and the area of each canopy gap was calculated. Then, canopy gap-related characteristics were calculated in a buffer of 60 m radius established around the centroid of each cell in the raster layer. The area of the buffer with such a radius was ∼1 ha, which was higher than the mean plus the standard deviation (SD) of the area of canopy gaps detected within the study area and contiguous areas (0.2 ha ±0.6 SD; range 0.03-4.96 ha); this allowed both small and large canopy gaps within the buffer to be included in the analyses. The canopy gap-related characteristics calculated for each cell buffer included the mean and SD of canopy gap areas (expressed in ha), the mean distance to the edge of canopy gaps (expressed in m), and the percentage of canopy gaps, i.e. the percentage of an area covered by canopy gaps. The calculations included all canopy gaps within or intersecting the buffer. To differentiate between cells with no canopy gaps within the buffer and cells whose centroids were in close proximity to or inside the canopy gaps (producing a mean distance = 0 m), we set a consistent mean distance of 85 m between the centroid and canopy gaps for cells without gaps within the buffer. Such a value corresponded to the mean plus SD of the distance between the ultimately selected adjoining canopy gaps (47.6 m ±36.5 SD, range 0.5-155.4 m), signifying the distance beyond which canopy gaps have a limited impact on the study plot. After this procedure, the mean distance from a cell’s centroid to the canopy gap represented a gradient from 0 to 85 m, i.e. from cells close to/inside canopy gaps to cells without canopy gaps.

Following previous studies on TreM prediction and modelling (Rehush et al., 2018; Frey et al., 2020; Santopuoli et al., 2020), stand structure metrics (Table S1) were calculated based on the raw LiDAR cloud of point measurements, which reflect the structure of a forest stand (Roussel et al., 2020). In the next step LiDAR-, topography- and canopy gap-related metrics were used to predict the density of living trees (for details, see Appendix 1), which was calculated for each cell in the raster layer.

### 2.4. Data handling and analyses

#### 2.4.1. Tree traits and TreM richness at the individual tree level

A total of 1683 trees – 1558 living trees and 125 snags – were examined during the fieldwork. As 98.9% were European beeches (Table S2), all analyses at the individual tree level were performed for European beeches (1547 living trees and 117 snags; Table S2). The basal area (the cross-sectional area of the tree trunk at breast height expressed in m^2^ ha^-1^) and TreM richness (expressed as total number of TreM types recorded on an individual tree) of each living tree and snag were calculated. The topographic characteristics, including elevation, slope, TPI, northness and eastness, and canopy gap-related metrics, i.e. the mean area of canopy gaps, the mean distance to the edge of canopy gaps and the percentage of canopy gaps, were read for each individual living tree and snag from the cell of the multi-layer raster (see Remote-sensed data section) where the tree/snag was located.

The mean TreM richness was calculated for all living trees and all snags pooled, and for all living trees and all snags separately. Then, the mean diameter, mean height and the percentage of individuals in 10 cm-width diameter classes were calculated. Thereafter, the number of trees in the diameter ranges defined on the basis of the increasing lower limit of the range in 10 cm-width intervals (i.e. 7.0-150 cm, 10.1-150 cm, 20.1-150 cm, 30.1-150 cm, …, 120.1-150.0 cm), the percentage of trees in a given diameter range and the percentage of TreM types recorded on trees in a given diameter range were calculated for all living trees and all snags separately. The differences in mean diameter, mean height and mean TreM richness between living trees and snags were tested using Student’s t-test. The multimodal distribution of the trees in the diameter classes was tested with Hartigans’ Dip Test for Unimodality (Hartigan and Hartigan, 1985).

Prior to the modelling procedures, the explanatory variables were tested for collinearity using Pearson’s correlation, and variable pairs with a correlation r ≤ 0.55 were included in the analyses (Fig. S1a-c). To address overdispersion and zero-inflation of model residuals in modelling count-type data, the relationships between the individual characteristics and TreM richness found on a given tree were analysed using Generalized Linear Mixed Models (GLMMs) with zero-inflated negative binomial error distribution and a log link function (Yirga et al. 2020). The set of 9 explanatory continuous variables, i.e. diameter, mean area of canopy gaps, mean distance to canopy gaps, percentage of canopy gaps, elevation, slope, TPI, northness and eastness, and the living status (living tree *vs* snag) as the categorical variable, were selected for the global model. Then, to reduce the number of variables, models with all possible combinations of all the variables from the global model were constructed, and the one with the lowest Akaike Information Criterion, corrected for a small sample size (AICc) (hereafter, best model), was selected. The categorical variables, i.e. the observer (two levels) and the identification numbers of the study plots (90 levels; from 1 to 90), were added as random effects to the best model to counteract the observer effect (Paillet et al., 2015) and the overdispersion of residuals, respectively. Zero-inflation of the residuals was counteracted by including the zero-inflation component in the best model, with the diameter and living status as explanatory variables, and the observer as a random effect. The AICc of the final model, i.e. the best model with random effects and a zero-inflation component, was lower than the global model (ΔAICc = 971.6). The residuals of the final model exhibited neither dispersion (p = 1.00; simulation-based dispersion tests) nor zero-inflation (p = 0.50; zero-inflation test).

#### 2.4.2. Habitat characteristics and TreM indices at the study plot level

All the analyses at the study plot level were based on the whole dataset, i.e. all 1683 trees (Table S2). A set of characteristics describing the forest stand structure and composition, topography, canopy gaps and TreM assemblage was calculated for each plot, and the mean, SD and range of these characteristics were calculated for all 90 study plots. The forest stand structure characteristics included the density, basal area and mean height of trees calculated separately for living trees and snags, while the forest stand composition characteristic was the number of tree species calculated on the basis of living trees and snags pooled. The topographic characteristics included elevation, slope, TPI, northness and eastness, and the canopy gap characteristics included the mean area of canopy gaps, the SD of the canopy gap area, the mean distance to canopy gaps and the percentage of canopy gaps. For each study plot, 40 metrics describing the cloud of LiDAR point measurements (Table S1) and predicted density of living trees (see Appendix 1) were calculated. The topography, canopy gap-related metrics, LiDAR metrics and predicted density of living trees were read from the cell of the multi-layer raster (see Remote-sensed data section), where the centre of the study plot was located.

The characteristics of the TreM assemblage were calculated on the basis of both living trees and snags pooled and included three TreM indices: TreM richness, i.e. the total number of TreM types recorded on a study plot; TreM density, i.e. the total density of trees bearing TreMs (if a tree bore several TreM types, it was counted once for each TreM type) extrapolated to 1 ha (expressed as TreM-bearing trees ha^-1^) (Paillet et al., 2017), and TreM diversity, i.e. the Shannon-Wiener diversity index calculated on the basis of TreM types recorded and their relative abundance (the density of trees bearing a given TreM type in the density of all TreM-bearing trees recorded on a study plot) (Przepióra and Ciach, 2022).

In accordance with the typology of Kraus et al. (2016), we grouped all 64 TreM types into twenty TreM groups and eight TreM forms. We calculated the density of TreM-bearing trees, i.e. the density of trees bearing a given TreM extrapolated to 1 ha, and frequency, i.e. the percentage of study plots with a given TreM, for each type, group and form. If particular TreM groups/forms were represented by several TreM types/groups on one tree, we counted the presence of a group/form only once. Therefore, the calculations of density and frequency of given TreM groups or forms on an individual tree were based on the occurrence of at least one TreM type from a given group or form, respectively.

The relationships between habitat characteristics and the three TreM indices found on a given study plot were analysed using GLMMs with Poisson error distribution and log link function in the case of TreM richness, and GLMMs with Gaussian error distribution and an identity link function in the case of TreM density and TreM diversity. The set of 14 continuous explanatory variables included the number of tree species, density of living trees, density of snags, basal area of living trees, basal area of snags, mean height of living trees, mean area of canopy gaps, mean distance to canopy gaps, percentage of canopy gaps, elevation, slope, TPI, northness and eastness. They were selected in order to build global models, after which the best model was selected (following the same procedure as with the models at the individual tree level). The final models were obtained by adding the categorical variable, i.e. the observer as a random effect to every best model for TreM richness, TreM density and TreM diversity. The AICc of each final model was lower than the corresponding global model, that is, for the TreM richness model (ΔAICc = 17.9), TreM density model (ΔAICc = 21.2) and TreM diversity model (ΔAICc = 21.9). The residuals of the final models exhibited overdispersion for TreM richness (p = 0.01) but not for TreM density (p = 0.50) or TreM diversity (p = 0.92).

#### 2.4.3. Spatial modelling and describing the spatial pattern of TreM indices

The TreM indices were predicted spatially across the investigated area using GLMMs with Gaussian error distribution and an identity link function in the case of TreM richness and TreM diversity, and GLMMs with gamma error distribution and log link function for TreM density. The set of explanatory variables used to build global models for the spatial prediction included 16 continuous variables: predicted density of living trees, elevation, slope inclination, TPI, northness and eastness, mean area of canopy gaps, SD of canopy gap area, mean distance to canopy gaps, percentage of canopy gaps, mean (ZMEAN), SD (ZSD) and entropy (ZENTROPY) of a point’s height, percentage of returns above mean height (PZABOVE1), 10^th^ percentile of height distribution of points (ZQ10), and percentage of 1^st^ returns (P1ST) in the LiDAR point cloud data (see Table S1). The set also included one categorical variable – the observer – treated as a random effect. Then, to reduce the number of variables, the models with all possible combinations of all variables from the global models were built and the models with the highest coefficient of determination (R^2^) selected.

The R^2^, Root Mean Square Error (RMSE), Normalized Root Mean Square Error (NMRSE) and uncertainty of each prediction were calculated using the leave-one-out cross-validation procedure. The uncertainty of prediction (U) was calculated as the range between the upper and lower 95% confidence intervals of prediction (CI) (Malone et al., 2017), normalized by the mean value of predictions (x) (Zhou et al., 2019) to allow comparison of the uncertainty parameters between the predictions of TreM indices. The following equation was used to calculate the uncertainty:

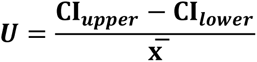

The final model was characterized by R^2^ = 0.26, RMSE = 4.20 and NRMSE = 0.19 for TreM richness (Table S3a, Fig. S2a), R^2^ = 0.23, RMSE = 287.72 and NRMSE = 0.21 for TreM density (Table S3b, Fig. S2b), and R^2^ = 0.28, RMSE = 0.10 and NRMSE = 0.19 for TreM diversity (Table S3c, Fig. S2c). The mean U was 0.32 ±0.09 SD (range 0.25-1.10) for TreM richness (Fig. S2d), 1.05 ±0.09 SD (range 0.49-1.53) for TreM density (Fig. S2e) and 0.14 ±0.09 SD (range 0.12-0.32) for TreM diversity (Fig. S2f). The predictions of TreM richness, density and diversity using the final models were carried out on the raster grid with the calculated explanatory variables (raster/grid of 1176 cells; see Remote-sensed data section). The prediction accuracy (R^2^ ∼0.30) was similar to the accuracy obtained in a previous study that attempted to predict the richness and abundance of TreMs using topography, canopy gap-related characteristics and laser scanning data (Frey et al., 2020).

The mean, skewness and histograms of distribution, that is, the frequency distributions of data, of the predicted TreM indices were calculated, and the normality of distribution was tested using the Shapiro-Wilk normality test. The spatial autocorrelations of the predicted TreM richness, density and diversity were tested using Moran’s I test (Li et al., 2007), and the spatial correlations between TreM richness, TreM density and TreM diversity on the raster grid (N = 1176) were tested using a modified t-test (Vallejos et al., 2020).

The predicted TreM indices for each individual grid cell were inspected using the Getis-Ord Gi* test, known as hot-spot analysis (Sussman et al., 2019). The test indicates whether the grid cells with high or low values are spatially clustered. It also calculates the difference between the sum of each grid cell and its neighbours, and the sum of all grid cells is significant (p < 0.05) if the difference is too large to be the result of random chance. A significant negative (z < 0) or positive (z > 0) z-score, which is the statistics of the test, indicates whether a grid cell belongs to a cluster of grid cells with relatively high values, i.e. a hot-spot, or a cluster of grid cells with relatively low values, i.e. a cold-spot, respectively. The grid cells with a non-significant (p > 0.05) z-score do not belong to any cluster, either a cold- or a hot-spot, so such grid cells are ranked as background.

The percentage of the area of grid cells ranked as cold-spot, hot-spot or background in the total area of all grid cells, and the mean area (expressed in ha) of individual clusters of grid cells ranked as cold- or hot-spot, were calculated. The individual clusters, i.e. individual cold- or hot-spots, were distinguished for all cells (with a resolution of 22.36 m) with the same rank connected by at least one edge. Differences between the mean areas of cold- and hot-spots were tested using the U Mann-Whitney test.

#### 2.4.4. Spatial visualization, software and packages

Three-dimensional (3D) spatial maps were created to visualize the study area, the spatial pattern of both TreM indices and cold- and hot-spots of TreM indices. The maps were created by generating a 3D layout based on the DEM, overlaying it with a grid of the predicted variables, both with a resolution of 0.5 m. To facilitate this, the resolution of the grid for which the prediction was made was changed from 22.36 m to 0.5 m, i.e. the corresponding value from the cell of the larger grid was assigned to each cell in the 0.5 m grid. All analyses were performed in R 3.5.0 software (CoreTeam R, 2017). Detection of canopy gaps, computation of topography variables and LiDAR variable calculations with tree top prediction (see Appendix 1, for details) were carried out using the ForestGapR (Silva et al., 2019), terra (Hijmans et al., 2022), and lidR (Roussel et al., 2020) packages, respectively. SpatialPack (Osorio et al., 2016) and multimode (Ameijeiras-Alonso et al., 2021) packages were employed for running modified t-tests and Hartigan’s Dip Test for Unimodality, respectively. GLMMs were run using the glmmTMB (Magnusson et al., 2017) package, and model diagnostics were conducted using the DHARMa (Hartig and Hartig, 2017) package. Moran’s I and the Getis-Ord Gi* tests were carried out using the Spdep (Bivand et al., 2015) and rgeoda (Li and Anselin, 2021) packages, respectively. For 3D visualizations, the rayshader (Morgan-Wall, 2023) package was employed.

## 3. Results

### 3.1. Tree traits and TreM richness at the individual tree level

The mean diameter of living trees was smaller (t = 2.47; p = 0.015) than the mean diameter of snags (Table 1a). The mean height of living trees was higher (t = −25.28; p < 0.001) than the mean height of snags (Table 1a). Distributions of living trees (p = 0.010) and snags (p < 0.001) were non-unimodal, in both cases with large contributions of individuals from the 10.1-20.0 cm diameter class and individuals from the > 60 cm diameter classes (Fig. S3). The mean TreM richness found on living trees and snags pooled was 2.0 ±3.4 SD TreM types per tree. The mean TreM richness found on living trees was 1.8 ±3.3 SD TreM types per tree and was significantly lower (t = 8.74; p < 0.001) than the mean TreM richness found on snags (4.7 ±3.5 SD TreM types per tree; Table 1a).

**Table 1.** (a) Mean ±SD and range of diameter at breast height, tree height and Tree-related Microhabitat (TreM) richness (total number of TreM types recorded on an individual tree); **(b)** density of trees, basal area of trees and mean height of trees, calculated separately for living trees and dead standing trees (snags); **(c)** TreM richness (total number of TreM types recorded on a study plot), TreM density (density of TreM-bearing trees ha^-1^), and TreM diversity (Shannon-Wiener index), calculated on the basis of living trees and snags pooled, found on the study plots; **(d)** elevation, slope inclination, Topographic Position Index (TPI), northness, eastness; and **(e)** mean area of canopy gaps, mean distance to canopy gaps and percentage of canopy gaps, calculated for the centre of study plots in the primeval European beech *Fagus sylvatica*-dominated forest in the Bieszczady Mountains (Poland).

| Characteristic | Unit | Living trees |  |  | Snags |  |  |
| --- | --- | --- | --- | --- | --- | --- | --- |
|  |  | Mean | ±SD | Range | Mean | ±SD | Range |
| (a) Individual tree traits |  |  |  |  |  |  |  |
| Diameter at breast height | cm | 31.2 | 24.8 | 7.0-150.0 | 37.9 | 28.7 | 7.0-125.0 |
| Tree height | m | 17.8 | 7.9 | 1.3-39.0 | 6.2 | 4.5 | 1.3-30.8 |
| TreM richness | TreM types per tree | 1.8 | 3.3 | 0.0-23.0 | 4.7 | 3.5 | 0.0-16.0 |
| (b) Stand structure |  |  |  |  |  |  |  |
| Density of trees | trees·ha<sup>-1</sup> | 346.2 | 164.7 | 120.0-880.0 | 27.8 | 31.1 | 0.0-200.0 |
| Basal area | m<sup>2</sup>·ha<sup>-1</sup> | 44.6 | 13.4 | 11.1-75.5 | 4.8 | 5.9 | 0.0-24.5 |
| Mean height of trees | m | 18.5 | 3.6 | 10.2-29.9 | 6.5 | 4.2 | 2.0-30.8 |
| (c) Tree microhabitats (TreMs) at the plot level |  | Living trees and snags pooled |  |  |  |  |  |
|  |  | Mean |  | ±SD |  | Range |  |
| TreM richness | TreM types per plot | 19.7 |  | 4.9 |  | 9.0-31.0 |  |
| TreM density | TreM-bearing trees·ha<sup>-1</sup> | 740.7 |  | 292.5 |  | 180.0-1520.0 |  |
| TreM diversity | Shannon-Wiener index | 1.2 |  | 0.1 |  | 0.9-1.4 |  |
| (d) Topography |  | Study plot centre |  |  |  |  |  |
| Elevation | m amsl | 921.2 |  | 51.5 |  | 822.8-1029.1 |  |
| Slope inclination | degrees | 21.0 |  | 7.0 |  | 7.2-41.1 |  |
| TPI |  | 0.1 |  | 1.4 |  | -3.7-3.0 |  |
| Northness | value from -1 to 1 | -0.6 |  | 0.4 |  | -1.0-0.7 |  |
| Eastness | value from -1 to 1 | 0.1 |  | 0.7 |  | -1.0-1.0 |  |
| (e) Canopy gaps |  |  |  |  |  |  |  |
| Mean area of canopy gaps | ha | 0.3 |  | 0.7 |  | 0.0-4.3 |  |
| Mean distance to canopy gaps | m | 35.1 |  | 14.2 |  | 0.0-85.0 |  |
| Percentage of canopy gaps | % | 10.7 |  | 7.5 |  | 0.0-40.2 |  |

The TreM richness of an individual tree was correlated with its living status: it was higher in snags (Table 2a, Fig. 2a), increased with diameter but decreased with the eastness of the slope where the tree was located (Table 2a). The TreM richness on an individual tree doubled with approximately every 30 cm of diameter growth in both living trees (Fig. 2b) and snags (Fig. 2c), and living trees had the same TreM type richness as snags with a 20 cm diameter range lag. The percentage of TreM types found on living trees or snags within a specific range of diameters decreased with the increasing lower limit of the range (Fig. 2d-e). Living trees in the 60.1-150 cm diameter range, i.e. 16.0% of all the living trees measured, provided 93.2% of all TreM types, while trees in the 120.1-150 cm diameter range, i.e. 0.4% of all the living trees measured, provided 47.5% of all TreM types (Fig. 2d). Snags in the 40.1-125 cm diameter range, i.e. 41.9% of all measured snags, and snags in the 90.1-125 cm diameter range, i.e. 3.4% of all measured snags, provided 90.9% and 50.0% of all TreM types found on all snags, respectively (Fig. 2e).

**Fig. 2.**
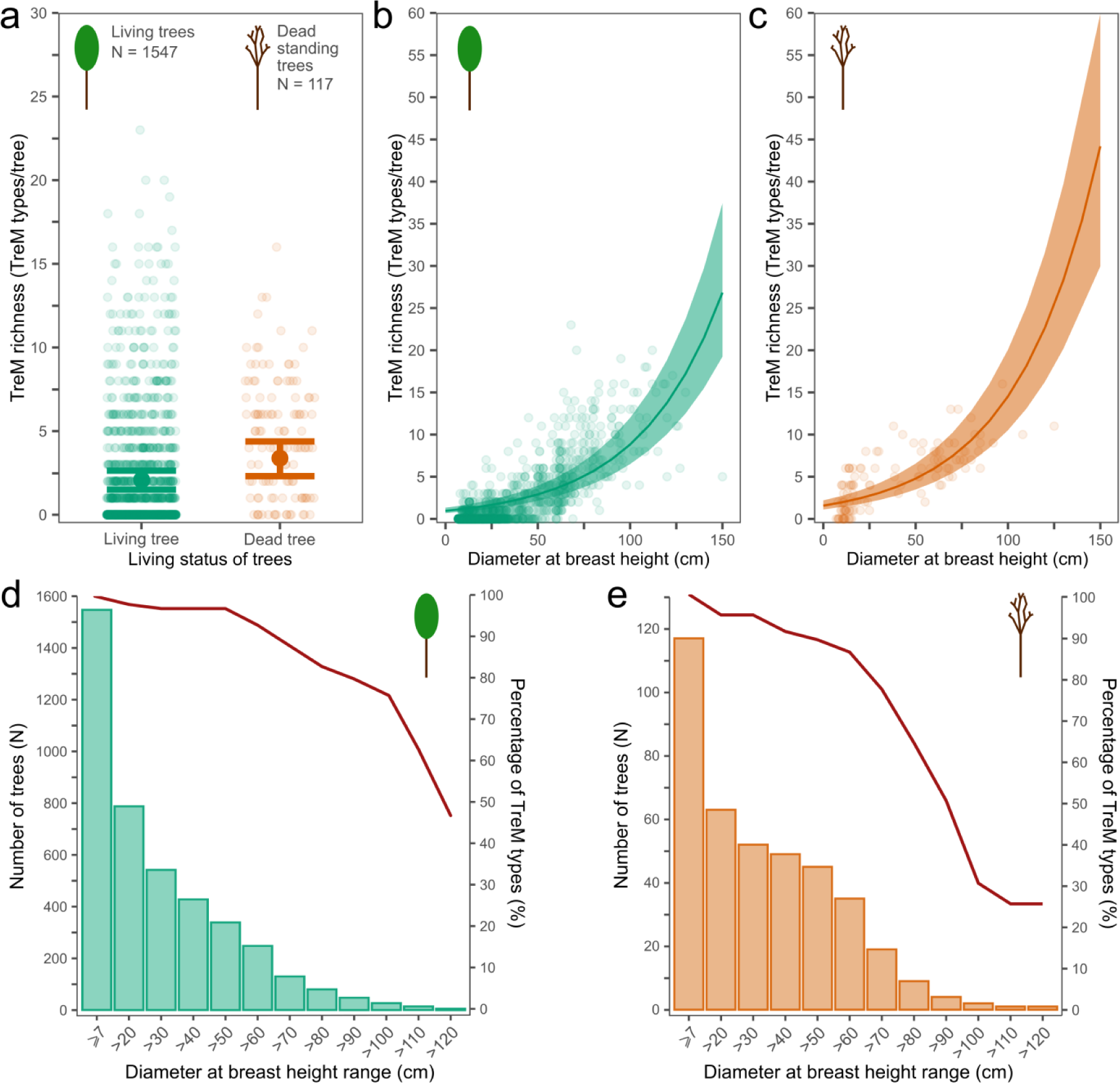
(a) Tree-related Microhabitat (TreM) richness (total number of TreM types recorded on an individual tree) found on living and dead standing trees; the relationships between TreM richness and diameter at breast height of **(b)** individual living or **(c)** dead standing trees; the relationships between the number of trees in a specific diameter range (bars; left axis) and percentage of TreM types (lines; right axis) calculated as the number of TreM types found on trees within a specific diameter range divided by the number of TreM types found on all trees based on measured **(d)** living trees and **(e)** dead standing trees in the primeval European beech *Fagus sylvatica*-dominated forest in the Bieszczady Mountains (Poland). Means – filled circles (a) or lines (b, c), and confidence intervals – whiskers (a) or ribbons (b, c) are products of the generalized linear-mixed models (Table 3a); the open circles are recorded data. The legend for the colours and sample sizes is shown inside panel a.

**Table 2.**
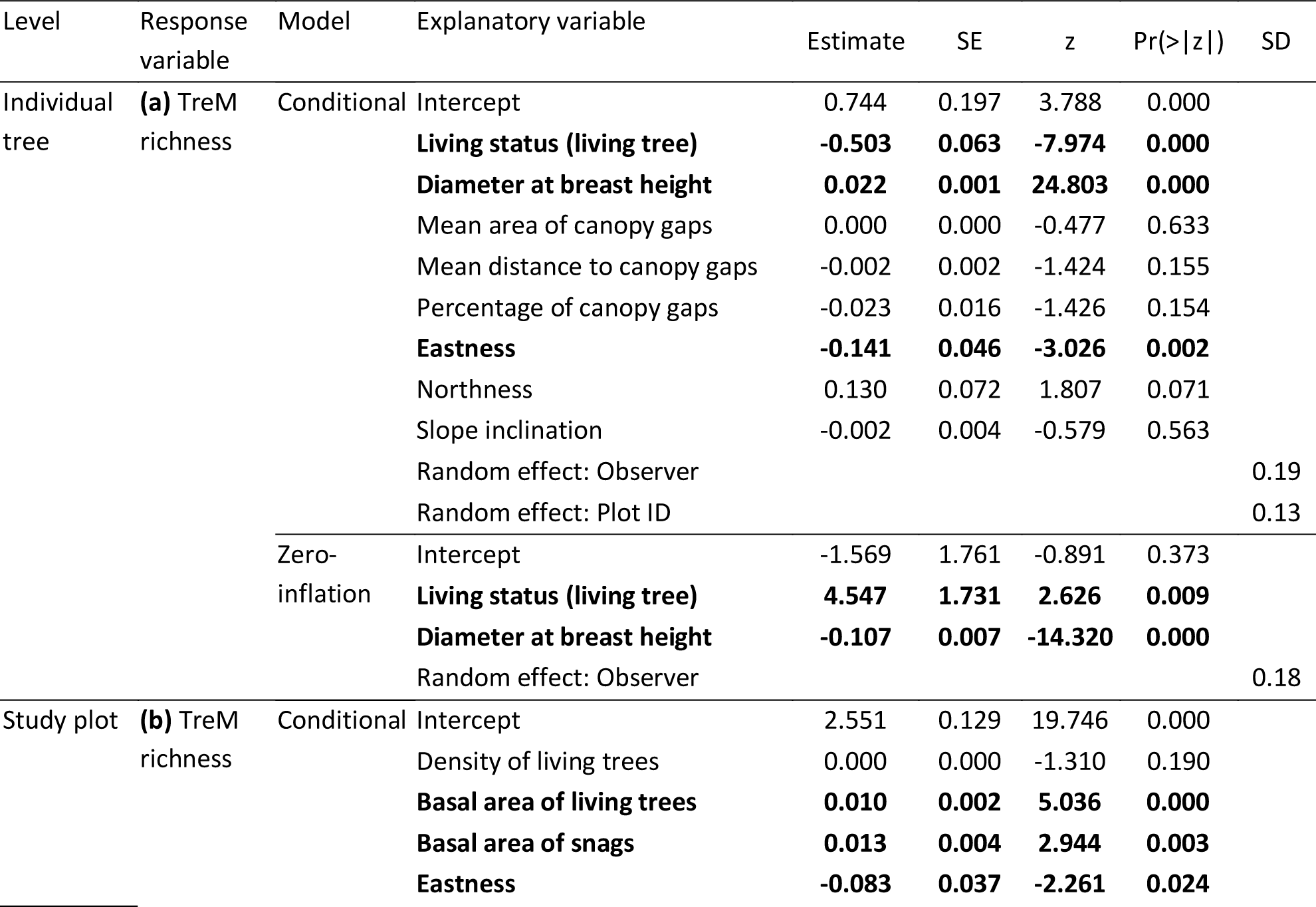
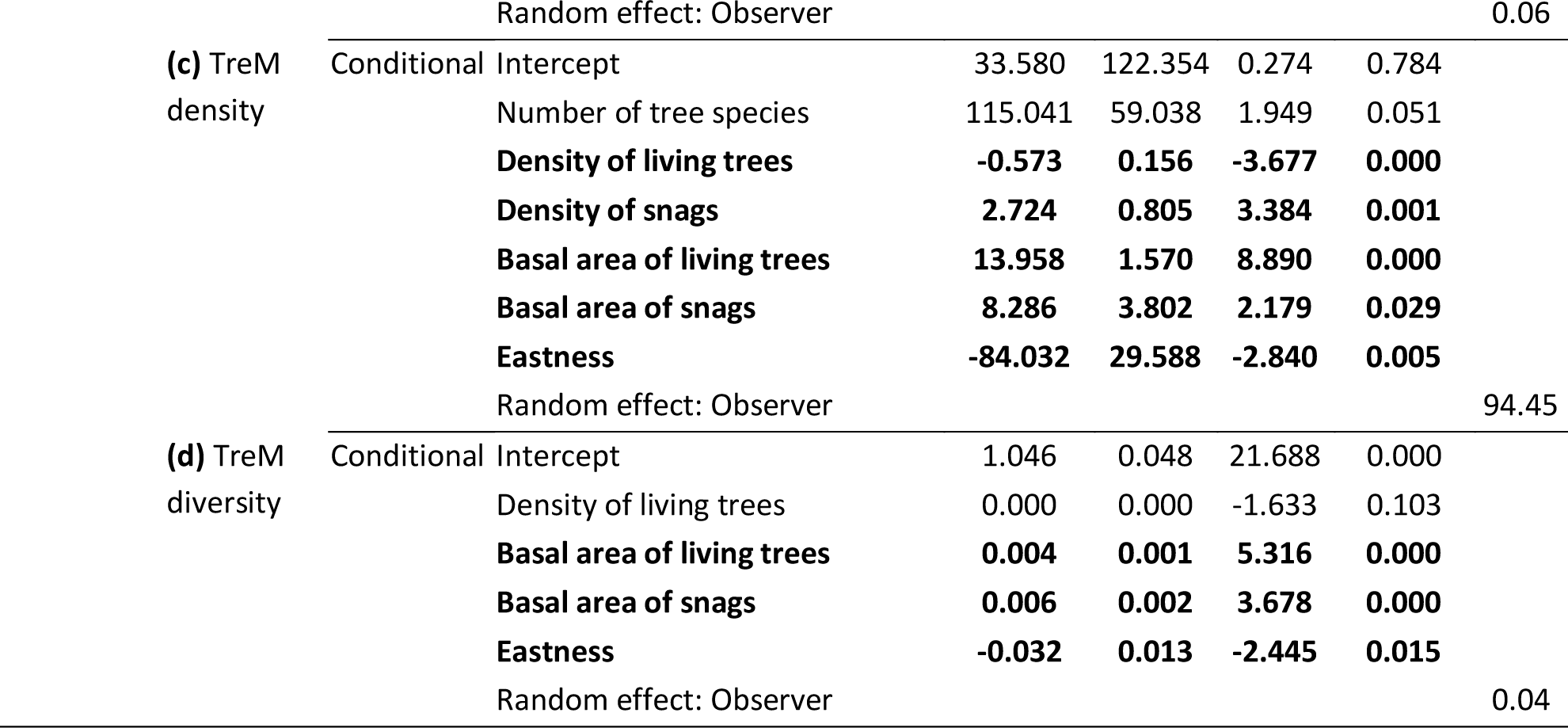
Models describing the relationships between **(a)** Tree-related Microhabitat (TreM) richness at the individual tree level (the total number of TreM types recorded on an individual tree) and living status (living *vs* dead standing tree), diameter at breast height, mean area of canopy gaps, mean distance to canopy gaps, percentage of canopy gaps, northness, eastness and inclination of slope (N = 1664); **(b)** TreM richness at the plot level (total number of TreM types recorded on a study plot); **(c)** TreM density (density of TreM-bearing trees ha^-1^); and **(d)** TreM diversity (Shannon-Wiener index) recorded on study plots in the primeval European beech *Fagus sylvatica-*dominated forest in the Bieszczady Mountains (Poland), and habitat characteristics (N = 90): number of tree species, density of living trees, density of snags (dead standing trees), basal area of living trees, basal area of snags and eastness of slope. The random effects, observer (two levels) and study plot ID (90 levels), and the zero-inflation model were used to counteract the observer effect, overdispersion of residuals and inflation of residuals, respectively. Significant results (Pr(>|z|) < 0.05) are in bold.

**Table 3.**
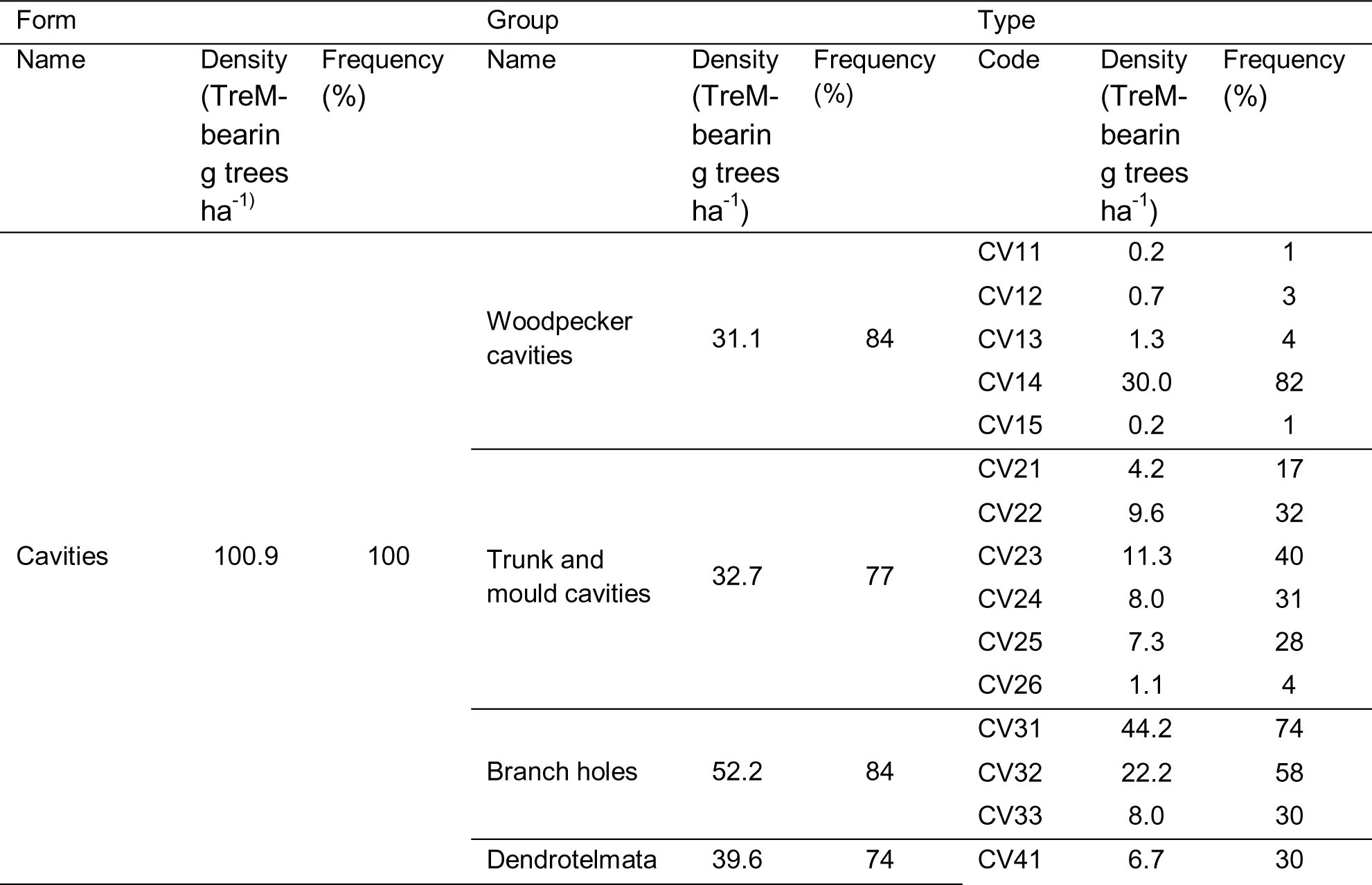
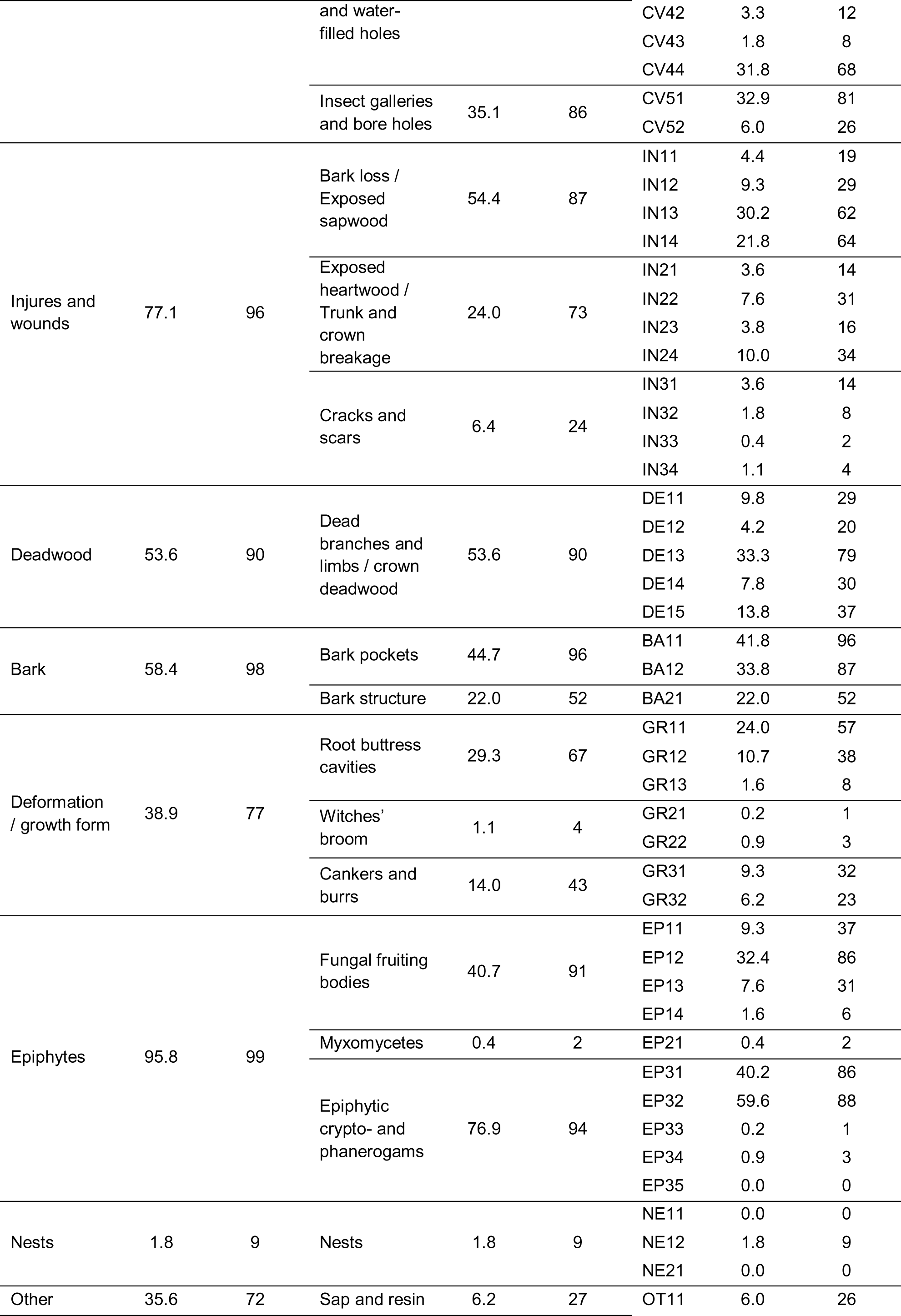
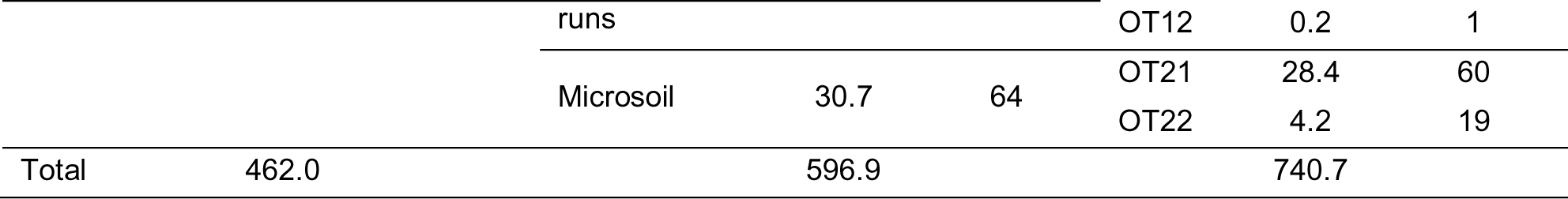
Tree-related Microhabitat (TreM) densities (density of trees bearing a given TreM ha^-1^) and frequencies (percentage of study plots with a given TreM, N = 90) of types, groups and forms of TreMs found in the primeval European beech *Fagus sylvatica*-dominated forest in the Bieszczady Mountains (Poland). The names of the TreM types, codes, groups and forms have been adapted from Kraus et al. (2016).

### 3.2. TreM assemblage and the relationship between habitat traits and TreM indices at the study plot level

Six tree species were found in the primeval European beech-dominated forest (Table S2), and the mean number of species found on the study plots was 1.1 ±0.4 SD (range 1.0-3.0). The mean density of living trees in the forest stand was 346.2 ±164.7 SD ha^-1^ with a mean basal area of 44.6 ±13.4 SD m^2^ ha^-1^ (Table 1b). The mean density and mean basal area of snags were 27.8 ±31.1 SD ha^-1^ and 4.8 ±5.9 SD m^2^ ha^-1^, respectively (Table 1b). The plots were characterized by a mean TreM richness of 19.7 ±4.9 SD TreM types per plot, a mean TreM density of 740.7 ±292.5 TreM-bearing trees ha^-1^, and a mean TreM diversity of 1.2 ±0.1 (Table 1c). The study plots were mostly situated on mountain slopes, usually with a south-eastern exposure (Table 1d). The mean size of the canopy gaps around the study plots was 0.3 ±0.7 SD ha, the mean distance between the centre of a study plot and the edges of canopy gaps was 35.1 ±14.2 SD m, and the mean percentage of areas assigned as canopy gaps around study plots was 10.7 ±7.5 SD % (Table 1e).

A total of 61 types of TreM were identified (Table 3). TreM types with the highest mean density were epiphytic foliose and fruticose lichens (TreM code EP32), small branch holes (CV31), bark shelter (BA11) and epiphytic bryophytes (EP31) (Table 3). TreM groups with the highest mean density were epiphytic crypto-and phanerogams, bark loss / exposed sapwood, dead branches and limbs / crown deadwood and branch holes (Table 3). TreM forms with the highest mean density were cavities and epiphytes (Table 3).

On the study plots, the TreM richness increased with the basal area of living trees and the basal area of snags, but decreased with the eastness of the slope where the study plot was located (Table 2b); the TreM density increased with the density of snags, basal area of living trees and the basal area of snags, but decreased with the density of living trees and slope eastness (Table 2c); the TreM diversity increased with the basal area of living trees and the basal area of snags, but decreased with the eastness of the slope (Table 2d).

### 3.3. Description of the spatial pattern of TreM indices

The cells with predicted TreM indices were not spatially distributed at random, indicating a clustered pattern (Moran’s I = 0.45-0.49; p < 0.001 for tests of each index) (Fig.3a-c). The frequency of cells with predicted TreM indices were not normally distributed (w = 0.91-0.99; p < 0.001 for tests of each index), and the skewness of the distribution was 0.25 for TreM richness, 1.49 for TreM density and 0.08 for TreM diversity, indicating right-skewed distributions (Fig. 3d-f). The cells with predicted TreM indices were characterized by a mean TreM richness of 20.8 ±1.8 SD TreM types per plot (Fig. 3d), a mean TreM density of 796.4 ±129.6 SD TreM-bearing trees ha^-1^ (Fig. 3e) and a mean TreM diversity of 1.3 ±0.1 SD (Fig. 3f). The spatial correlation was significant between each pair of TreM indices and was the highest between TreM richness and TreM diversity (r_XY_ = 0.93), followed by that between TreM diversity and TreM density (r_XY_ = 0.76), and that between TreM richness and TreM density (r_XY_ = 0.67) (Fig. 4).

**Fig. 3.**
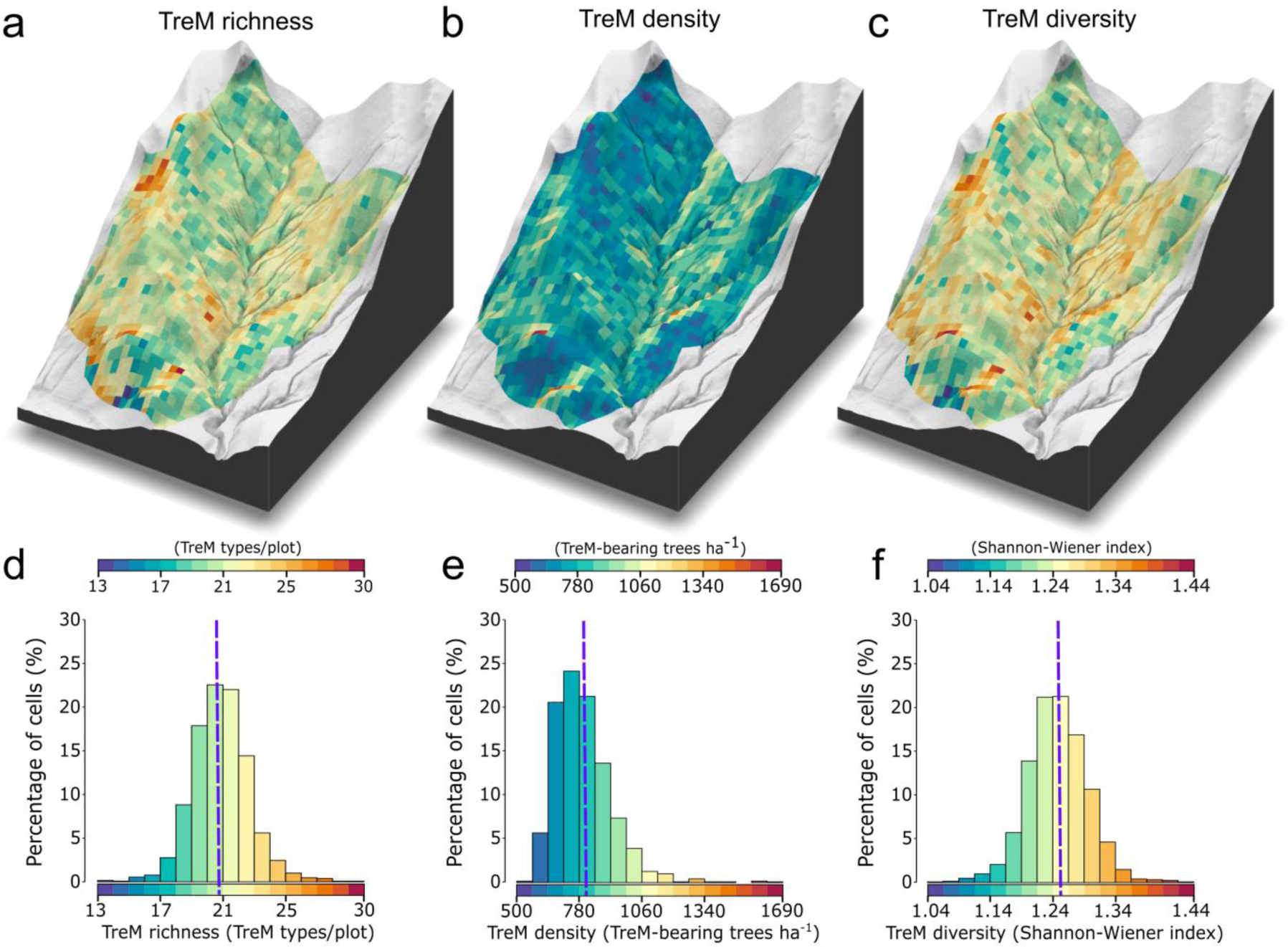
(a) Spatial distribution of predicted Tree-related Microhabitat (TreM) richness (total number of TreM types recorded on a plot), **(b)** TreM density (density of TreM-bearing trees ha^-1^) and **(c)** TreM diversity (Shannon-Wiener index) in the primeval European beech *Fagus sylvatica*-dominated forest in the Bieszczady Mountains (Poland), and the histograms of predicted **(d)** TreM richness, **(e)** TreM density and **(f)** TreM diversity expressed as a percentage of cells with a specific value. The colour legends for all the panels are shown at the top of panels d-f. The vertical purple dashed line indicates the mean value of the predicted TreM index calculated as the sum of all values from all cells divided by the total number of cells.

**Fig. 4.**
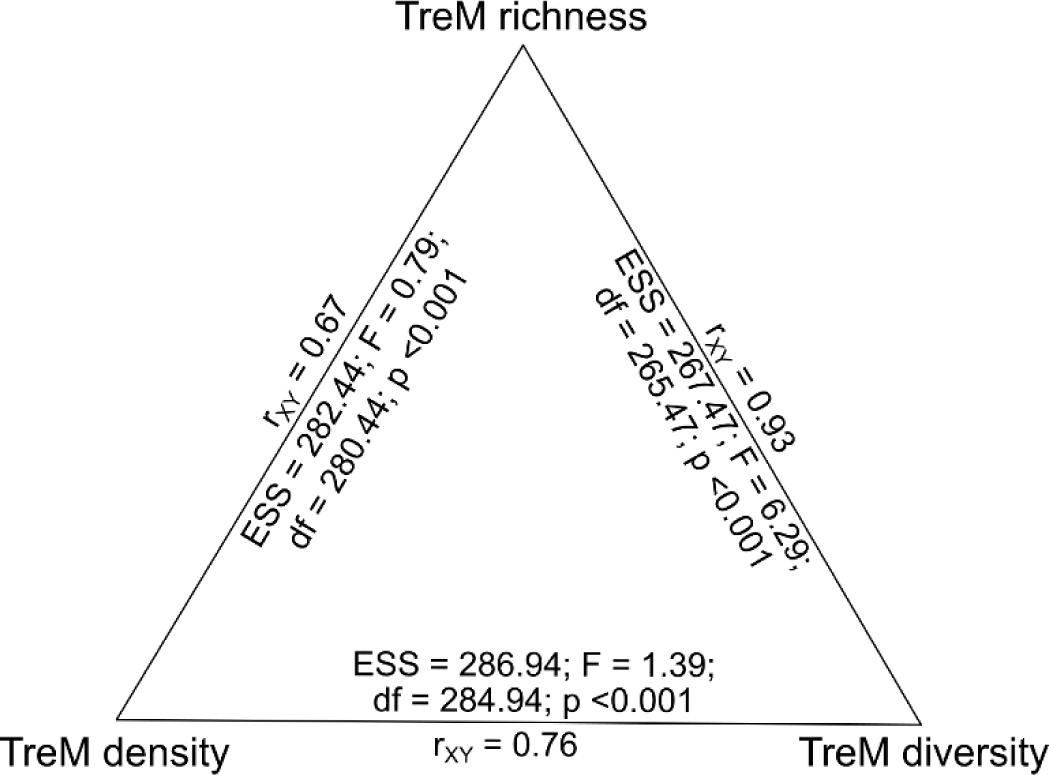
Correlation coefficient (r_XY_), effective sample size (EES), F statistic (F), degrees of freedom (df) and p value of the modified t-test (Vallejos et al., 2020) determining the measure of spatial correlation between predicted Tree-related Microhabitat (TreM) richness (total number of TreM types recorded on a study plot), TreM density (density of TreM-bearing trees ha^-1^) and TreM diversity (Shannon-Wiener index) in the primeval European beech *Fagus sylvatica*-dominated forest in the Bieszczady Mountains (Poland).

The percentages of cell areas ranked as cold- or hot-spots in the areas of all cells were similar for all the TreM indices. For TreM richness (Fig. 5a), the cells classified as cold-spots constituted 18.1% of the area of all cells, and the cells classified as hot-spots accounted for 17.9% (Fig. 5d). For TreM density (Fig. 5b), the cells classified as cold-spots made up 24.3% of the area of all cells, and the cells classified as hot-spots accounted for 19.2% (Fig. 5e). For TreM diversity (Fig. 5c), the cells classified as cold-spots constituted 17.5% of the area of all cells, and cells classified as hot-spots accounted for 22.2% (Fig. 5f). The mean areas of cold- or hot-spots of TreM richness were 0.66 ±0.27 SE ha (range 0.05-4.23) and 0.81 ±0.24 SE ha (range 0.05-2.54), respectively. In the case of TreM density, the mean areas of cold- or hot-spots were 0.90 ±0.53 SE ha (range 0.05-8.03) and 1.02 ±0.53 SE ha (range 0.05-6.17), respectively. The mean areas of TreM diversity cold- or hot-spots were 0.71 ±0.25 SE ha (range 0.05-2.91) and 1.05 ±0.51 SE ha (range 0.05-5.80), respectively. There were no differences in the mean sizes of cold- and hot-spots for all the TreM indices (Fig. 5g-i).

**Fig. 5.**
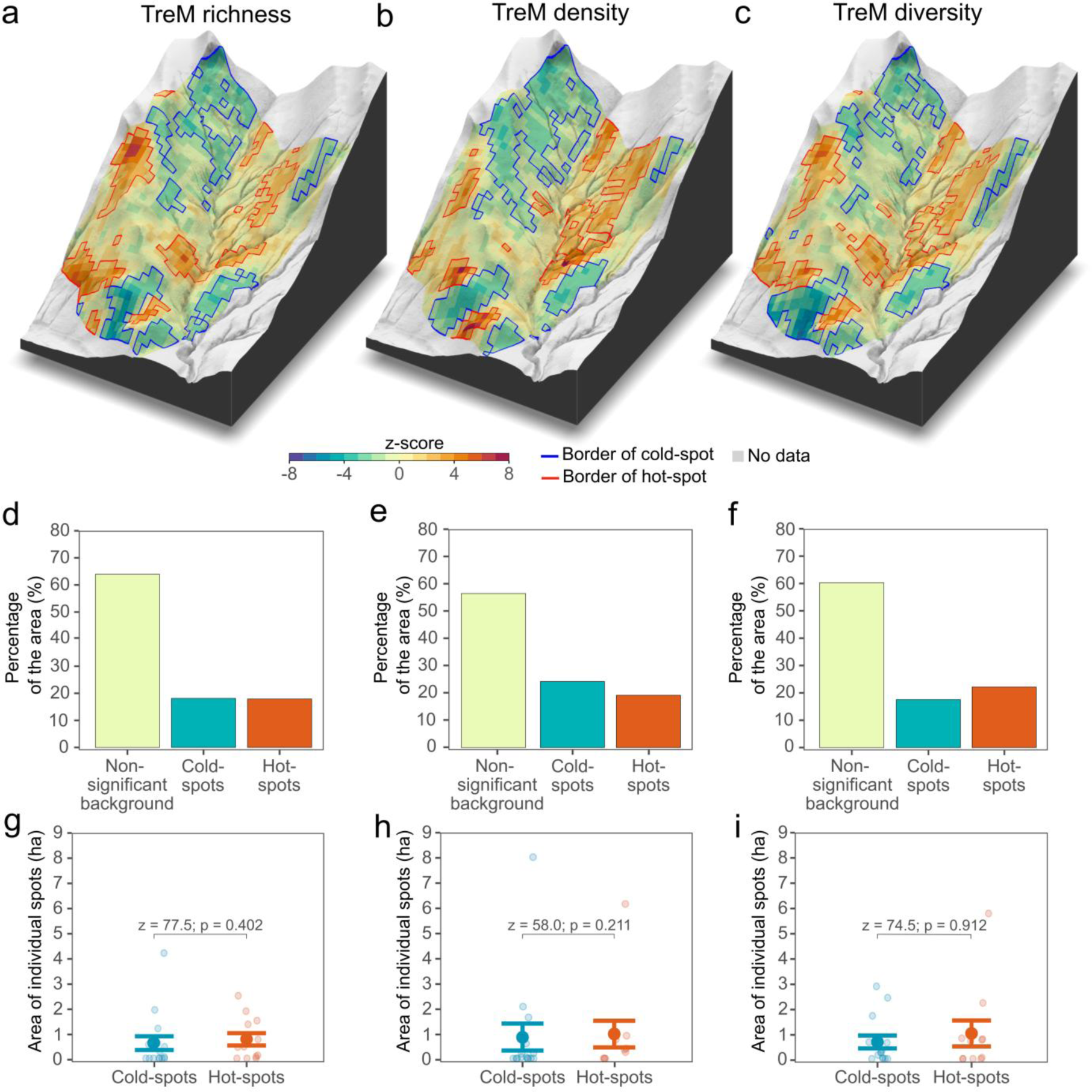
Spatial distribution of cold- and hot-spots (clusters of forest patches with significantly lower or higher indices in relation to the values of neighbouring forest patches) of predicted **(a)** Tree-related Microhabitat (TreM) richness (total number of TreM types recorded on a study plot), **(b)** TreM density (density of TreM-bearing trees ha^-1^) and **(c)** TreM diversity (Shannon-Wiener index) in the primeval European beech *Fagus sylvatica*-dominated forest in the Bieszczady Mountains (Poland); percentage of the area of cold- or hot-spots and non-significant background (forest patches which were neither cold- nor hot-spot) of **(d)** TreM richness, **(e)** TreM density and **(f)** TreM diversity in the total area of the forest and the mean (filled circles) ±SD (whiskers) of the area of cold- or hot-spots of **(g)** richness, **(h)** density and **(i)** TreM diversity.

## 4. Discussion

### 4.1. Profile and reference values of TreMs

Our study provides references for the quantity and quality of TreMs in primeval mountain European beech forests. While several studies conducted to date have examined TreMs in forests dominated by European beech (Winter and Möller, 2008) or mixed with other species (e.g. beech-fir forest; Larrieu and Cabanettes, 2012, oak *Quercus* sp.-beech-hornbeam *Carpinus* sp. forest; Paillet et al., 2017), those stands have been subject to long drawn out human transformations and are not considered primeval (Sabatini et al., 2018). Secondary or previously managed forests have very much smaller numbers of snags and large living trees, resulting in the absence of certain TreM types (Larrieu et al., 2012; Paillet et al., 2017). As a result, managed forests typically have fewer TreMs and do not have TreM profiles typical for ancient stands (Asbeck et al., 2022).

Unmanaged European beech forests in mountain regions like the Carpathians have persisted in spite of challenging forest management practices (UNESCO, 2021), and those remaining patches are rich in TreMs (Kozak et al., 2018; Sever and Nagel, 2019; Jahed et al., 2020). However, the TreM density in the Bieszczady Mts. is higher than that in other parts of the Carpathians. Specifically, the TreM group density in the forest we studied (596.9 TreM-bearing trees ha^-1^) was higher than that described in the Bukovské and Maramures Mts. (480.0 TreM-bearing trees ha^-1^; Kozák et al., 2018) and in the Uholka Forests (106.2 TreM-bearing trees ha^-1^; Jahed et al., 2020). Furthermore, we found that the density of certain types/groups of TreMs varied between the studied regions. For instance, the densities of woodpecker cavities (31.1 TreM-bearing trees ha^-1^) and fungal fruiting bodies (40.7 TreM-bearing trees ha^-1^) were 95.6% and 23.0% higher, respectively, than in the Bukovské and Maramures Mts. (15.9 and 33.1 TreM-bearing trees ha^-1^ for woodpecker cavities and conks of fungi, respectively; Kozák et al., 2018). In contrast, the densities of bryophytes (i.e. EP31 – 40.2

TreM-bearing trees ha^-1^ in our study, compared to 57.2 TreM-bearing trees ha^-1^ in Kozák et al., 2018) or cracks and scars (6.4 TreM-bearing trees ha^-1^ in our study, compared to 23.7 TreM-bearing trees ha^-1^ in Kozák et al., 2018 and 9.3 TreM-bearing trees ha^-1^ in Jahed et al., 2020) were lower than in other parts of the Carpathians. The reasons for the regional variation in TreM profiles in European beech-dominated forests is understudied, but could be the outcome of local climate, topography or disturbance regimes (Mráz and Ronikier, 2016; Kozak et al. 2018).

### 4.2. Effect of tree traits and stand structure on TreMs

We found that tree-level TreM richness increased with diameter and was higher in snags. TreM richness doubled for approximately every 30 cm of growth in diameter, and living trees achieved the same TreM richness as snags with a 20 cm diameter class lag. The majority of TreMs require a tree to have a long lifespan, during which it sustains various kinds of damage or wood decay as a result of the disturbances and activities of TreM-forming biota, with alternating wound healing and tissue superstructure (Kõrkjas et al., 2021). Moreover, a tree exposed for a long time to rare abiotic factors, such as lightning, can develop unique TreMs such as cracks with charred wood (Larrieu et al., 2018). Therefore, the TreM richness of an individual tree increases with trunk diameter (Kõrkjas et al., 2021), which in turn is strongly correlated with the tree’s age (Churski and Niklasson, 2010). After a tree’s death, the resulting snags host diverse TreMs following the formation of structures associated with wood decay, such as fungal fruiting bodies and patches of loose bark (Bütler et al., 2013).

Although snags host richer sets of TreMs than living trees (Paillet et al., 2019; Larrieu et al., 2022), the combined presence of both, especially those with larger diameters, leads to the most diverse stand-level TreM composition (Spînu et al., 2022; Przepióra and Ciach, 2023). The richness, density and diversity of TreMs we found increased with the basal area of both living trees and snags. The tree population we studied exhibited a bimodal distribution of diameter classes and included a significant proportion of large living trees and snags with diameters above 60 cm, the presence of which could have been responsible for a local increase in the basal area (Cade, 1997). Hence, the presence together of both large living trees and snags could have led to the high TreM indices we found in the European beech-dominated forest we studied.

In our study, living trees with diameters above 60 cm and 120 cm contained 93% and 47% of all the TreM types found on all the living trees, respectively. For European beech trees in mixed forests, the importance of trees as TreM hosts increases rapidly above a diameter of 70 cm (Larrieu et al., 2012; Großmann et al., 2018). Therefore, the harvesting of such large trees seriously depletes the TreM assemblage. For example, the elimination of trees with diameters larger than 50 cm in beech-fir forests reduces the number of TreMs by 48% and destroys one-sixth of all TreM types (Larrieu and Cabanettes, 2012). Hence, defining both a diameter threshold above which a tree should be retained and the number of such unlogged trees would be of practical value in landscape planning and forest management. In our study, trees with diameters > 60 cm reached a density of ca 55 trees per ha, which is in line with previous studies suggesting a minimum of 60 trees with diameters > 65 cm to ensure the continuing existence of a complex assemblage of TreMs in beech-fir forests (Larrieu et al., 2014). Therefore, we recommend retaining ≥ 55 living trees with diameters > 60 cm per ha in mountain European beech-dominated forests to preserve approximately 93% of the TreM types found in the assemblage typical of a primeval forest. Since the TreM richness, density, and diversity increased with the availability of snags, we emphasize the importance of retaining snags of any diameter because of their great value resulting from ongoing wood decomposition and the probable development of distinctive TreMs that rarely develop on living trees.

### 4.3. Effect of canopy gaps on TreMs

Our study failed to reveal any relationship between TreM indices and the presence of canopy gaps and their characteristics. A previous study of mixed Norway spruce-European beech-silver fir forests showed that the TreM density increased with the percentage of canopy gaps (Frey et al., 2020), although this research was conducted in a managed forest, where certain ecological processes, including TreM formation, may be disrupted as a result of former tree removal (Asbeck et al., 2022). In natural European beech forests, canopy gaps, originating mostly from small-scale disturbances caused by strong winds or snowfall, lead to the accumulation of TreM-rich dead, wounded and damaged living trees (Lewandowski et al., 2021). However, canopy gaps also offer increased sunlight exposure and lower humidity, which could negatively affect the occurrence of certain TreMs, like fungi, bryophytes or lichens (Larrieu et al., 2018; Esseen and Ekström, 2023) and retard the formation of mould cavities (Remm and Lõhmus, 2011). Therefore, canopy gaps may involve a trade-off between different types of TreMs on trees: some TreM types benefit from canopy gaps, while others have fewer chances of developing in such conditions. Consequently, the TreM profile on trees near canopy gaps may change, but the overall number of TreM types will remain fairly constant. Therefore, it is challenging to capture the relationship between canopy gaps and the TreM assemblage in heterogeneous old-growth forest using general TreM indices, due to the complex processes taking place within the ecosystem.

Canopy gaps may, however, have a delayed and indirect effect on the TreM assemblage, creating opportunities for the spontaneous growth of willows *Salix* spp., poplars *Populus* spp. or sycamore, which are TreM-rich pioneer or sun-demanding species (Przepióra and Ciach, 2023). Although such species accounted for just 1% of all the trees measured in the studied forest, the increasing number of tree species found on the study plots was close to being a significant predictor of the TreM-bearing tree density (see Table 2c). This suggests that the emergence of canopy gaps as a consequence of small-scale disturbances increases forest heterogeneity, initiating the appearance of additional tree species in the species pool, which in turn may boost TreM numbers in the mature stand. Hence, the emergence of canopy gaps, leading to the appearance of such long-distance wind-dispersed species or sun-demanding tree species should be supported in managed forests.

### 4.4. Effect of topographic conditions on TreMs

We observed a decrease in TreM richness on individual trees, as well as in the richness, density and diversity of TreMs on the study plots with increasing eastness of the slope. The study area is predominantly affected by southerly and south-westerly winds (Wibig, 2021), and hurricane-force winds from the west can cause considerable tree damage in Central European forests, including the Bieszczady Mts. (State Forests, 2018). Trees on west-facing slopes are more vulnerable to wind damage, giving rise to TreMs that include branch and trunk breakage. Wind also favours buttress formation, i.e. reaction wood that forms root buttress cavities (Crook et al., 1997). Furthermore, wind induces asymmetric tree crowns (MacFarlane and Kane, 2017), which are susceptible to snow damage, fairly common in beech forests (Homma, 1997), that increase the number of TreMs on trees growing on western slopes. However, Asbeck et al. (2020) found a greater TreM richness on eastern slopes, explaining such a pattern by the greater number of European beech trees growing in mixed forests on slopes with this aspect.

In the studied forest, there was also a nearly significant positive correlation between TreM richness on an individual tree and its position on north-facing slopes (see Table 2a). Slope exposure affects tree size and stand structure, potentially having an indirect influence on the number and diversity of TreMs at both the individual tree and stand levels. South-facing slopes in the Northern Hemisphere may receive significantly more solar radiation (Auslander et al., 2003), resulting in a warmer and drier microclimate that promotes taller and thicker European beech trees on north-facing slopes (Diaconu et al., 2015). North-facing slopes, with their more humid microclimate, favour tree cavity formation (Remm and Lõhmus, 2011) and epiphyte growth (Esseen and Ekström, 2023), potentially leading to an increased abundance of TreMs. However, the relationship between TreM numbers and north-facing slopes was not evident in mixed beech-spruce-fir forests (Asbeck et al., 2022). Hence, our findings suggest that the influence of slope exposure on TreMs depends on the local landscape context that determines the climatic conditions prevailing in the studied region.

We found no relationships between TreM numbers and slope inclination or elevation. A higher elevation may alter local rainfall and humidity, promoting tree cavity and epiphyte formation (Remm and Lõhmus, 2011; Kozák et al., 2023). Steep mountain slopes can be the cause of tree trunk damage due to rockfall (Šilhán, 2010), which leads to the development of reactive wood and asymmetrical root systems, and also the formation of root buttress cavities (Young and Perkocha, 1994). Such patterns were observed in fir-beech forests, as stands growing on steep slopes at high elevations host more TreMs, including buttress cavities and epiphytes (Asbeck et al., 2019). However, our study suggests that the local land configuration has a limited impact on TreMs, which, in turn, may be more evident at larger spatial scales, e.g. between mountain ranges.

### 4.5. Spatial distribution of TreMs

The spatial pattern of TreM indices in temperate forests has not been extensively investigated to date. Our study revealed the spatial pattern of TreM richness, density and diversity in a primeval mountain forest dominated by European beech. Previous research conducted in boreal forests identified variation in the distribution of habitat trees and TreM indices among forest patches of varying size, tree age and disturbance history (Martin et al., 2021a; Martin et al., 2021b). A study conducted in Oak-lime *Tilia* sp.-hornbeam forests revealed spatial variation in TreM indices and their dependence on the forest structure, i.e. tree density and basal area, and local variation in the tree species composition (Przepióra and Ciach, 2023).

Abiotic factors, such as relief, hydrological and soil conditions, and local disturbances, create diverse habitat patches that differ in the size and age of trees (Franklin and Van Pelt, 2004) and microclimatic conditions (Frey et al., 2016), leading to the formation of different local TreM assemblages over an extended period of time. Biotic factors, such as the activity of ecosystem engineers such as woodpeckers or bears and ungulates, further shape the occurrence of such TreMs as cavities or tree injuries, respectively (Bütler et al., 2004; Zyśk-Gorczyńska et al., 2015; Broughton et al., 2022). Moreover, TreMs such as drying-out dendrotelmata or the fruiting bodies of annual fungi are distinct ephemeral resource patches (Larrieu et al., 2018), i.e. they are only present for a limited period of time, which further considerably diversifies TreM occurrence.

We found a clustered pattern in the TreM richness, density and diversity distribution, where forest patches with low or high TreM indices formed cold- and hot-spots, respectively. Such cold- and hot-spots each occupied about 20% of the area, whereas the habitat background was characterized by an intermediate number of TreMs, covering about 60% of the forest. Our findings have implications for retention forestry, which aims to preserve key components of old-growth forests like TreM-rich patches. Depending on size, such retained habitat patches can serve as protected core areas, or those located between larger protected areas can create ecological corridors in the form of stepping-stone habitat patches (Vandekerkhove et al., 2013; Kuuluvainen et al., 2021). Larrieu et al. (2012) recommend excluding 10-20% of the forest area from management to ensure a high number of TreM-bearing trees at the landscape scale. Augustynczik et al. (2019) suggested protecting 10% of the area and reducing the volume of harvested trees by 10% on the remaining area in order to combat TreM decline due to climate change. Our study defines the proportion of an area that should be excluded from management in a system of protected/managed forests and sets up the ‘2:6:2’ rule, which allocates 20% to forests as strict protected areas, 60% to low-intensive forestry with reduced harvesting and the retention of large living trees and all snags, and the remaining 20% to timber production. Our recommendation is in line with the European Union’s 2030 Biodiversity Startegy, which urges member countries to increase the coverage of protected areas to 30%, with one-third under strict protection (Hermoso et al., 2022).

Our results also characterize the size of forest patches that should be designated for protection. We found that cold- or hot-spot patches averaged around 1 ha, with maximum sizes of 8 and 6 ha, respectively. This supports the concept of establishing a functional network of forest patches that act as stepping stones between forest reserves or national parks, with sizes ranging from 1-5 ha (Vandekerkhove et al., 2013). A network of such forest patches, called Woodland Key Habitats (WKHs), was previously established in boreal forests (Ylisirniö et al., 2016). It was shown that most WKHs, ranging from 0.5-5 ha, host at least one red-listed bryophyte or lichen species (Gustafsson et al., 1999). The proportion of WKHs with an average size of 2.5 ha in the landscape was also demonstrated to be the best predictor of species richness of red-listed insects (Götmark et al., 2011). The > 1 ha threshold size of WKH was advocated as sustaining the full polypore species diversity over the long term (Ylisirniö et al., 2016). Although no such threshold recommendations for the size of patches to be designated for protection in temperate deciduous forests have been drawn up, studies from the boreal zone suggest that habitat patches as outlined in our study may also be sufficient for the occurrence of some rare and endangered species.

However, it is important to note that in order to preserve all TreM types, along with the associated assemblages of organisms, larger areas of protected forests are necessary. It was shown that a minimum of 20 ha of unmanaged forest was necessary to provide the full assemblage of TreMs in beech-fir forests (Larrieu et al., 2014). Hence, in certain forest types, individual forest patches of the size examined in our study, i.e. 1-6 ha, may not guarantee the complete assemblage of TreMs or suitable habitats for some forest species. This is especially true for species with large area requirements. For example, white-backed woodpecker needs 50-100 ha of suitable habitat for breeding (Martikainen et al., 1998), while Eurasian lynx *Lynx lynx* needs forest complexes > 30 km^2^ in area for permanent residence, using smaller forest patches only for passage (Schadt et. al., 2002). We therefore recommend that such TreM-rich patches should be retained in addition to larger areas of old-growth forests. Moreover, while retaining habitat islands to mitigate the loss of old-growth forest, we also recommend the prioritization of patches with rich, abundant and diverse TreM assemblages rather than those dominated by a single TreM type, as patches with a high TreM density alone may lack TreMs crucial for a great many target species.

## 5. Conclusion

Our study revealed the density and frequency of particular TreMs and the richness, abundance and diversity of the TreM assemblage in a primeval temperate mountain European beech-dominated forest in the northern Carpathians. Since the studied forest is among the best-preserved, the values of the TreMs presented in this study can be considered as references and contribute to understanding the natural TreM occurrence pattern in beech forests. Our work shows that the local stand structure and conditions related to slope exposure, but not to the presence of canopy gaps, influence the number and diversity of TreMs in a given forest patch. We also revealed the spatial pattern of TreM indices, which are spatially diversified and clustered in TreM-rich and TreM-poor patches. Such TreM hot- and cold-spots each covered approximately 20% of the forest and were situated within a forest matrix with average TreM indices. Based on our findings, we advocate the ‘2:6:2’ rule, namely, that 20% of forests should be allocated as strict protected areas, 60% dedicated to low-intensive forestry with reduced harvesting and the retention of large living trees and all snags, and the remaining 20% used for intensive timber production. In low-intensity managed forests, a minimum of 55 living trees per ha with a diameter > 60 cm should be preserved in order to save the majority of TreM types. As TreM-rich patches have low densities of living trees, high densities of snags and high basal areas of both living trees and snags, forest managers can rely on existing forest structure data when designating habitat patches for protection. Our results contribute to the concept of triad forestry, which aims to mitigate the negative effects of forest management on biodiversity.

## Acknowledgements

This study was financially supported by The National Science Centre in Poland from a Preludium grant (2021/41/N/NZ9/03441) and an Opus grant (2021/41/B/NZ8/03456), and also by the Ministry of Science and Higher Education of the Republic of Poland within the framework of statutory funds awarded to the Faculty of Forestry, University of Agriculture. We thank the management of the Bieszczady National Park for permission to carry out this study and Stanisław Kucharzyk for his consultancy during the selection of the study sites. This work is dedicated to the memory of the late Jerzy Lesiński, advocate of forest management methods that comply with the need for nature conservation.

## Ethical statement

The study was performed in accordance with Polish law.

## Supplementary Materials

**Table S1.** Mean ±SD and range of metrics describing Light Detection and Ranging point measurements taken within the boundaries of the investigated area of the primeval European beech *Fagus sylvatica*-dominated forest in the Bieszczady Mountains (Poland).

| Variable | Description and unit | Mean | SD | Range |
| --- | --- | --- | --- | --- |
| ZMAX | Height of the highest point [m] | 30.6 | 4.4 | 18.1-40.5 |
| ZMEAN | Mean height of point [m] | 15.4 | 3.5 | 6.8-24.3 |
| ZSD | Standard deviation of height of points [m] | 9.3 | 1.8 | 5.3-13.0 |
| ZSKEW | Skewness of height distribution of points | -0.5 | 0.5 | -2.0-1.2 |
| ZKURT | Kurtosis of height distribution of points | 2.1 | 0.8 | 1.2-6.4 |
| ZENTROPY | Entropy of height of points | 0.9 | 0.0 | 0.7-1.0 |
| PZABOVE1 | Percentage of returns above mean height of points [%] | 58.8 | 6.5 | 34.2-73 |
| PZABOVE2 | Percentage of returns above 2 [%] | 80.6 | 7.7 | 58.4-91.9 |
| ZQ10 | 10<sup>th</sup> percentile (quantile) of height distribution of points [%] | 0.3 | 1.2 | 0.0-10.0 |
| ZQ15 | 15<sup>th</sup> percentile (quantile) of height distribution of points [%] | 2.5 | 3.8 | 0.0-20.2 |
| ZQ20 | 20<sup>th</sup> percentile (quantile) of height distribution of points [%] | 4.8 | 5.0 | 0.0-22.2 |
| ZQ25 | 25<sup>th</sup> percentile (quantile) of height distribution of points [%] | 7.6 | 5.6 | 0.0-23.4 |
| ZQ30 | 30<sup>th</sup> percentile (quantile) of height distribution of points [%] | 10.1 | 5.7 | 0.0-24.3 |
| ZQ35 | 35<sup>th</sup> percentile (quantile) of height distribution of points [%] | 12.4 | 5.5 | 0.5-25.1 |
| ZQ40 | 40<sup>th</sup> percentile (quantile) of height distribution of points [%] | 14.4 | 5.3 | 1.3-26.0 |
| ZQ45 | 45<sup>th</sup> percentile (quantile) of height distribution of points [%] | 16.1 | 5.1 | 3.9-26.9 |
| ZQ50 | 50<sup>th</sup> percentile (quantile) of height distribution of points [%] | 17.6 | 4.8 | 5.8-27.6 |
| ZQ55 | 55<sup>th</sup> percentile (quantile) of height distribution of points [%] | 18.9 | 4.6 | 7.0-28.2 |
| ZQ60 | 60<sup>th</sup> percentile (quantile) of height distribution of points [%] | 20.0 | 4.6 | 8.6-28.8 |
| ZQ65 | 65<sup>th</sup> percentile (quantile) of height distribution of points [%] | 21.0 | 4.6 | 9.8-29.7 |
| ZQ70 | 70<sup>th</sup> percentile (quantile) of height distribution of points [%] | 22.0 | 4.5 | 10.8-30.6 |
| ZQ75 | 75<sup>th</sup> percentile (quantile) of height distribution of points [%] | 22.9 | 4.5 | 11.7-31.5 |
| ZQ80 | 80<sup>th</sup> percentile (quantile) of height distribution of points [%] | 23.9 | 4.5 | 12.5-32.2 |
| ZQ85 | 85<sup>th</sup> percentile (quantile) of height distribution of points [%] | 24.8 | 4.5 | 13.0-33.0 |
| ZQ90 | 90<sup>th</sup> percentile (quantile) of height distribution of points [%] | 25.9 | 4.5 | 13.7-33.8 |
| ZQ95 | 95<sup>th</sup> percentile (quantile) of height distribution of points [%] | 27.2 | 4.5 | 14.7-34.6 |
| ZPCUM1 | Cumulative percentage of return in the 1<sup>st</sup> layer [%] | 9.1 | 6.8 | 1.1-34.1 |
| ZPCUM2 | Cumulative percentage of return in the 2<sup>nd</sup> layer [%] | 13.7 | 9.7 | 2.1-50.4 |
| ZPCUM3 | Cumulative percentage of return in the 3<sup>rd</sup> layer [%] | 18.2 | 11.6 | 2.5-61.5 |
| ZPCUM4 | Cumulative percentage of return in the 4<sup>th</sup> layer [%] | 23.9 | 12.9 | 3.3-80.4 |
| ZPCUM5 | Cumulative percentage of return in the 5<sup>th</sup> layer [%] | 32.2 | 13.8 | 4.6-85.0 |
| ZPCUM6 | Cumulative percentage of return in the 6<sup>th</sup> layer [%] | 43.9 | 14.6 | 10.6-85.5 |
| ZPCUM7 | Cumulative percentage of return in the 7<sup>th</sup> layer [%] | 59.7 | 14.3 | 24.8-88.0 |
| ZPCUM8 | Cumulative percentage of return in the 8<sup>th</sup> layer [%] | 78.2 | 10.5 | 49.3-96.3 |
| ZPCUM9 | Cumulative percentage of return in the 9<sup>th</sup> layer [%] | 94.4 | 3.9 | 79.1-99.7 |
| P1ST | Percentage of 1<sup>st</sup> returns [%] | 51.4 | 4.9 | 41.8-63.6 |
| P2ND | Percentage of 2<sup>nd</sup> returns [%] | 33.0 | 1.5 | 28.1-36.0 |
| P3RD | Percentage of 3<sup>rd</sup> returns [%] | 12.6 | 2.5 | 6.6-18.2 |
| P4TH | Percentage of 4<sup>th</sup> returns [%] | 2.7 | 1.2 | 0.3-5.5 |
| P5TH | Percentage of 5<sup>th</sup> returns [%] | 0.3 | 0.2 | 0.0-1.1 |

**Table S2.** Numbers of sampled living trees and dead standing trees (snags) of European beech *Fagus sylvatica*, silver birch *Betula pendula*, goat willow *Salix caprea*, sycamore *Acer pseudoplatanus*, European aspen *Populus tremula*, Norway spruce *Picea abies* and all species pooled in the primeval mountain European beech-dominated forest in the Bieszczady Mountains (Poland).

| Tree species | Living trees | Snags | Pooled (living trees and snags) |
| --- | --- | --- | --- |
| All species pooled | 1558 | 125 | 1683 |
| European beech *Fagus sylvatica* | 1547 | 117 | 1664 |
| Silver birch *Betula pendula* | 2 | 2 | 4 |
| Goat willow *Salix caprea* | 3 | 4 | 7 |
| Sycamore *Acer pseudoplatanus* | 2 | 0 | 2 |
| European aspen *Populus tremula* | 1 | 0 | 1 |
| Norway spruce *Picea abies* | 3 | 2 | 5 |

**Table S3.**
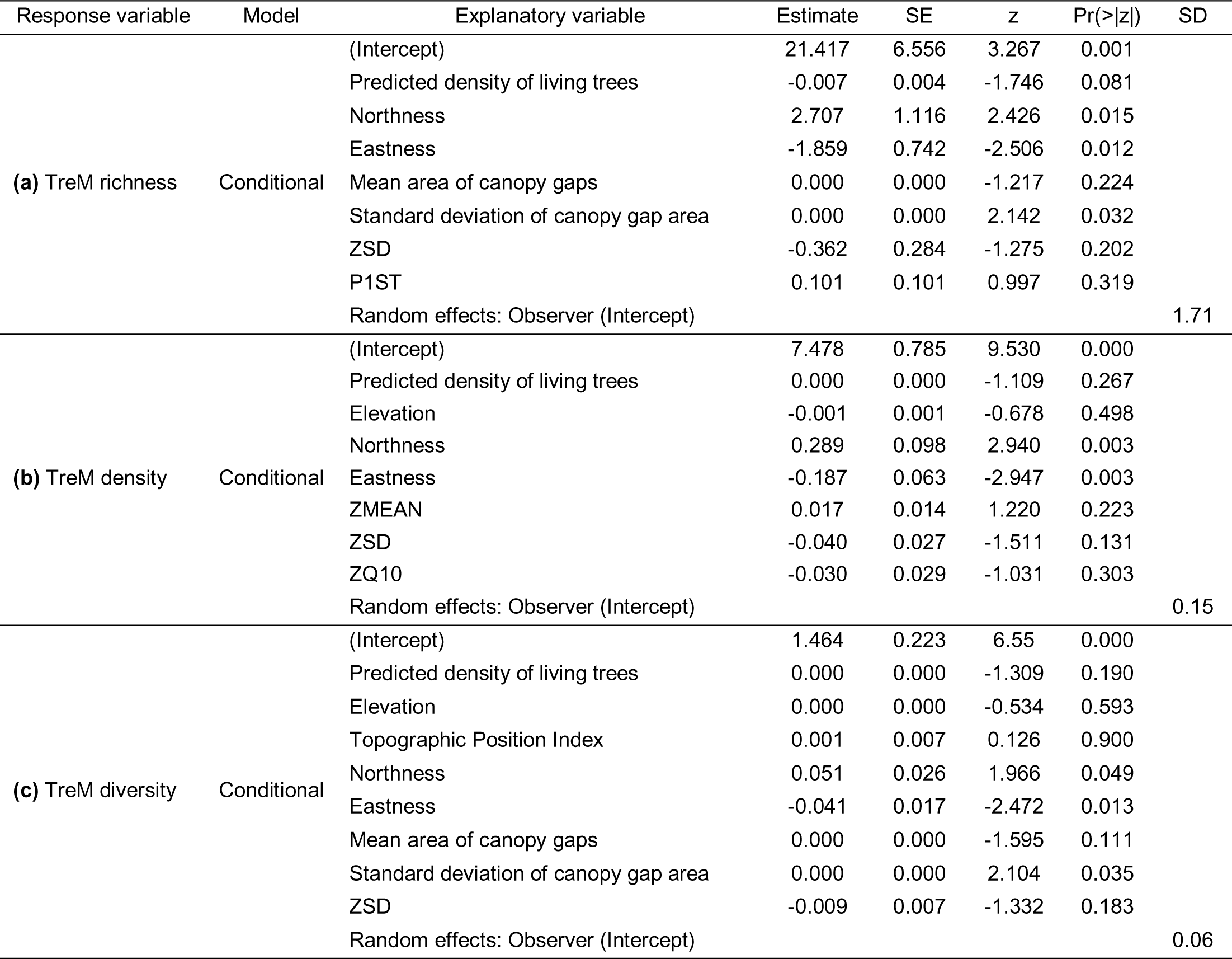
Final models used to predict **(a)** Tree-related Microhabitat (TreM) richness (total number of TreM types recorded on a study plot), **(b)** TreM density (density of TreM-bearing trees ha^-1^) and **(c)** TreM diversity (Shannon-Wiener index) within the investigated area of the primeval European beech *Fagus sylvatica*-dominated forest in the Bieszczady Mountains (Poland) based on remote data: predicted density of living trees (see Appendix 1), elevation, Topographic Position Index, northness, eastness, mean area of canopy gaps, standard deviation of canopy gap area, ZMEAN (mean pixel height), ZSD (standard deviation of pixel height), P1ST (percentage of 1^st^ returns), ZQ10 (10^th^ percentile (quantile) of height distribution of points). The random effect - observer (two levels) was used to account for the observer effect (Paillet et al., 2015).

**Fig. S1.**
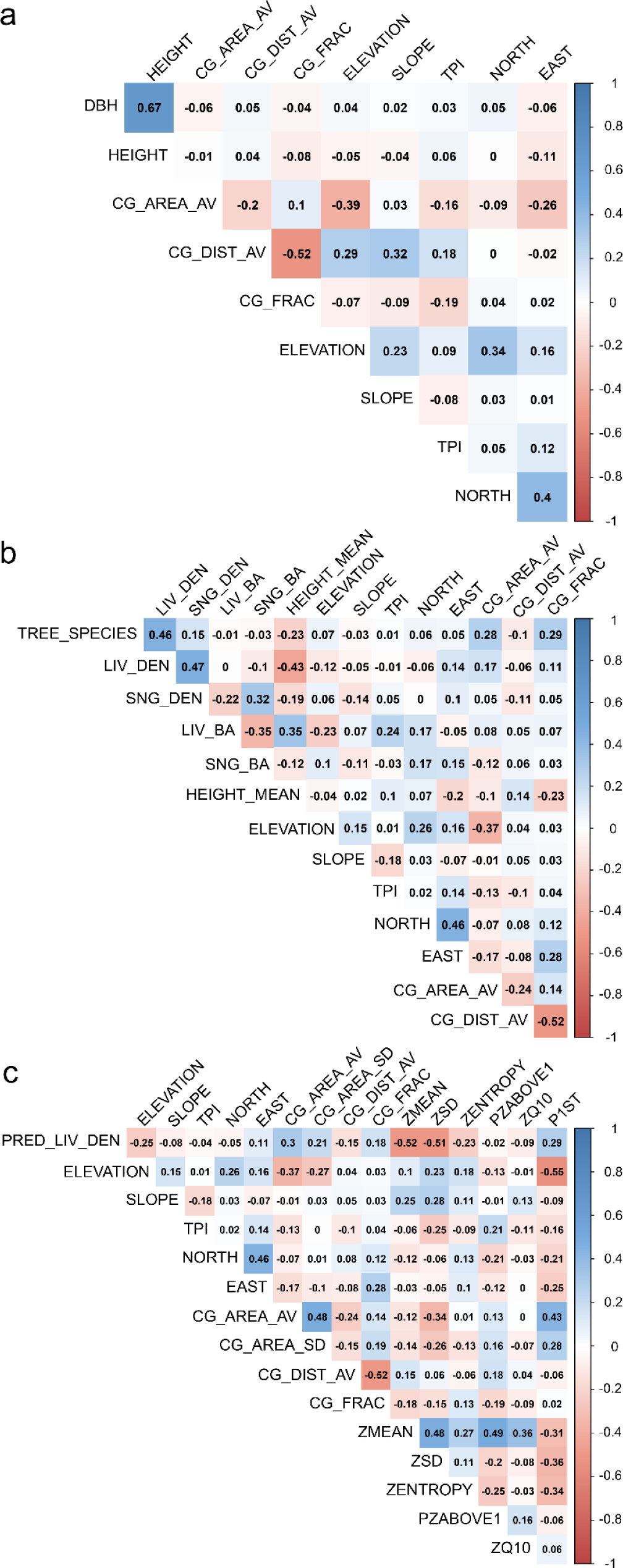
Pearson correlation coefficients between tree and habitat characteristics: diameter at breast height (DBH), tree height (HEIGHT), mean canopy gap area (CG_AREA_AV), standard deviation (SD) of canopy gap area (CG_AREA_SD), mean distance to canopy gaps (CG_DIST_AV), percentage of canopy gaps (CG_FRAC), elevation (ELEVATION), slope inclination (SLOPE), Topographic Position Index (TPI), northness (NORTH), eastness (EAST), number of tree species (TREE_SPECIES), density of living trees (LIV_DEN), density of dead standing trees (SNG_DEN), basal area of living trees (LIV_BA), basal area of dead standing trees (SNG_DEN), height of living trees (HEIGHT_MEAN), predicted density of living trees (PRED_LIV_DEN), mean height of Light Detection and Ranging (LiDAR) points (ZMEAN), SD of height of LiDAR points (ZSD), entropy of height of LiDAR points (ZENTROPY), percentage of returns above ZMEAN (PZABOVE1), 10^th^ percentile (quantile) of height distribution of LiDAR points (ZQ10) and percentage of 1^st^ returns in LiDAR point cloud (P1ST) used as continuous explanatory variables in models describing relationships between **(a)** Tree-related Microhabitats (TreM) and individual tree traits, **(b)** TreMs and study plot characteristics and **(c)** TreMs and spatial-available characteristics of the primeval European beech *Fagus sylvatica*-dominated forest in the Bieszczady Mountains (Poland). Because of the large number of variables, panel c shows only those with correlations significant at the adopted r ≤ 0.55.

**Fig. S2.**
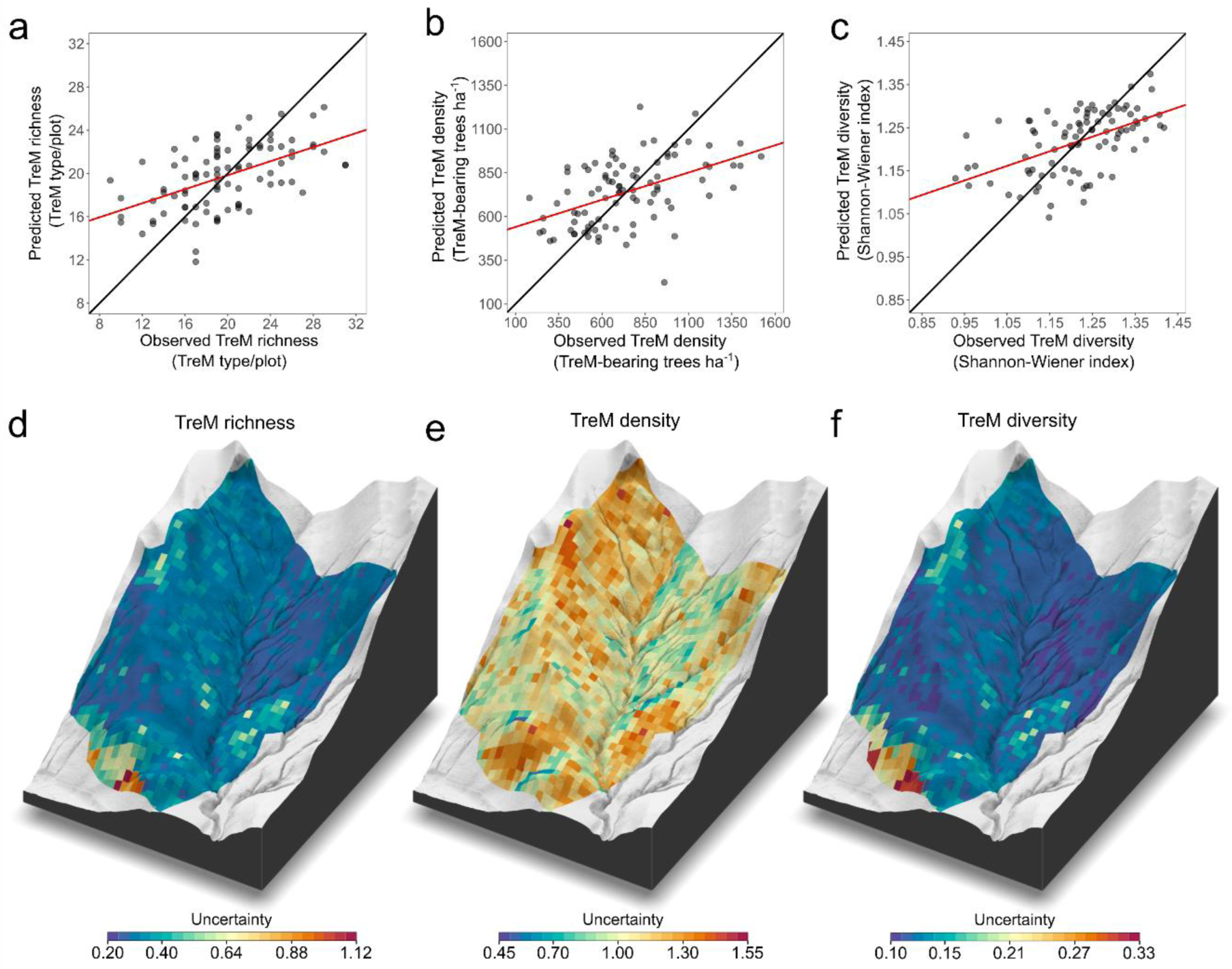
The relationship between predicted and observed **(a)** Tree-related Microhabitat (TreM) richness (total number of TreM types recorded on a study plot), **(b)** TreM density (density of TreM-bearing trees ha^-1^) and **(c)** TreM diversity (Shannon-Wiener index) found on study plots in the primeval European beech *Fagus sylvatica*-dominated forest located in the Bieszczady Mountains (Poland), and the spatial distribution of uncertainty of prediction of **(d)** TreM richness, **(e)** TreM density and **(f)** TreM diversity. The black lines in panels a-c indicate theoretical linear relationships with a slope equal to 1 and the red lines indicate the calculated linear relationship between the observed and predicted TreM indices.

**Fig. S3.**
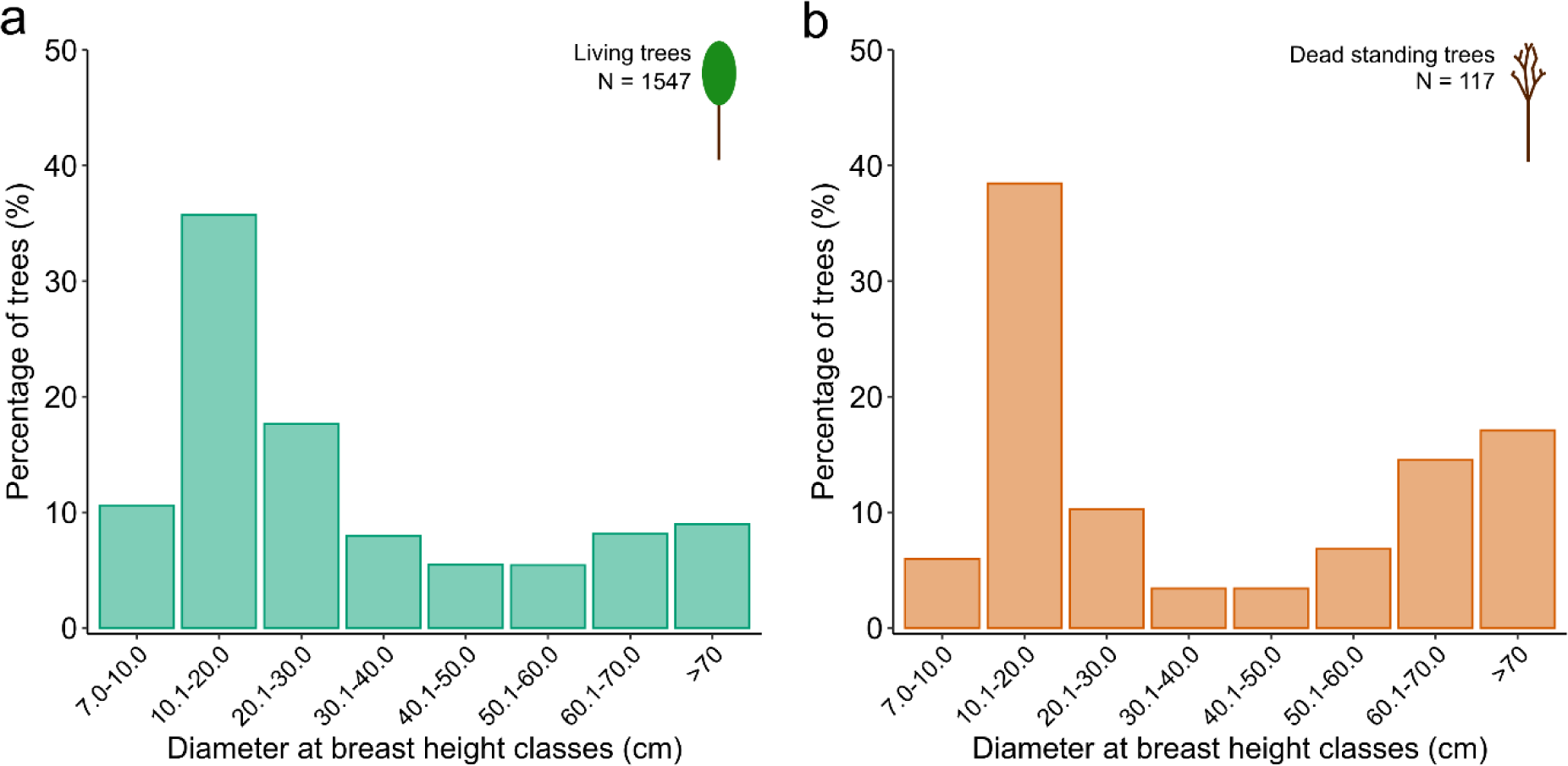
Distribution of **(a)** living trees and **(b)** dead standing trees of European beech *Fagus sylvatica* in diameter at breast height classes (cm) in the primeval European beech-dominated forest in the Bieszczady Mountains (Poland).

## Appendix 1

### Remote data – prediction of density of living trees

The predicted living tree densities were based on the treetops of dominant trees, which were detected using fixed-window-size local-maxima (LM) filtering (Popescu and Wynne, 2004; Gebreslasie et al., 2011) applied to the Canopy Height Model (CHM) acquired from Light Detection and Ranging (LiDAR) data (obtained from airborne laser scanning with a density of 4 points per m^2^; https://www.geoportal.gov.pl/en/data/lidar-measurements-lidar). This method is based on the assumption that a treetop is the highest point within a crown and that the crown boundary is relatively low (Zhen et al., 2016; Xu et al., 2021). A pixel in CHM is identified as an LM when its neighbouring pixels have a lower height value (Koch et al., 2006; Plakman et al., 2020). The LM filter is processed across CHM using a sliding circular or a square search window of specific size (Xu et al., 2021). A large window is suitable for large scattered trees with extensive crowns whereas a small window is preferable for small and closely-adjacent trees with relatively smaller crowns (Douss and Farah, 2022). Therefore, in more complex and heterogenous stands, in which the trees are of varying size and age structure, the search window size should be adjusted to one that corresponds to the tree crowns (Popescu and Wynne, 2004; Ke and Quackenbush, 2011; Zhen et al., 2016; Xu et al., 2021).

The size of the search window in this method is a function of the height of the detected treetop. The relationship between crown size and tree height is species-specific (Popescu and Wynne, 2004; Buchacher and Ledermann, 2020). The function usually used in the LM method is a linear one whose coefficients are fitted to a particular species or stand type (e.g. Pitkänen et al., 2004; Chen et al., 2020; Plakman et al., 2020). To date, few papers have described a function between tree height and search window size in complex, old hardwood forests. In our study, therefore, the values of the coefficients in the function were adjusted to match the search window size to conditions in the European-beech *Fagus sylvatica*-dominated forest stand we studied. The matching process involved checking combinations of different values of the coefficients and the minimum and maximum window sizes. For each combination, we detected treetops and then evaluated the coefficient of determination (R^2^) between the number of detected treetops and the number of living trees measured within the boundaries of the study plots. The final function formula with the highest accuracy of treetop detection (R^2^ = 0.34), which described the relationship between the diameter of the circular search window (ws) and the height of the LM pixel (H) in the CHM we used (Fig. A1), was as follows:

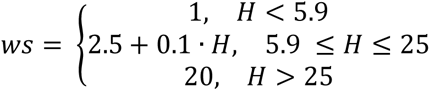

After the best function had been chosen, the detection of treetops over the whole study area was performed using the lidR package (CoreTeam R., 2017; Roussel et al., 2020). Then, the entire area was covered with a vector grid with a resolution of 22.36 m, and the numbers of detected treetops within each grid cell and within the boundaries of each study plot were calculated.

The spatial prediction of the living tree density across the investigated area was performed using Generalized Linear Mixed Models with Gaussian error distribution and an identity link function. The set of explanatory variables used for the spatial prediction included the number of detected treetops and topography characteristics, canopy gap characteristics and 40 LiDAR metrics (see Remote data section). Prior to the analysis, all the explanatory continuous variables had been tested for collinearity using Pearson’s correlation, and the variable pairs with correlation r > 0.55 were excluded. The set of 16 explanatory continuous variables were selected to build the global model; they included the number of detected treetops, elevation, slope inclination, Topographic Position Index, northness, eastness, mean area of canopy gaps, SD of the canopy gap area, mean distance to canopy gaps, percentage of canopy gaps and the mean (ZMEAN), SD (ZSD) and entropy (ZENTROPY) of point heights, the percentage of returns above the mean height (PZABOVE1), the 10^th^ percentile of the height distribution of points (ZQ10) and the percentage of 1^st^ returns (P1ST) in the LiDAR point cloud data (see Table S1). Then, to reduce the number of variables, the models with all possible combinations of all variables from the global model were built and the best model with the lowest Akaike Information Criterion corrected for a small sample size and the highest R^2^ of the prediction were selected. The R^2^, Root Mean Square Error (RMSE) and Normalized Root Mean Square Error (NRMSE) of each prediction were calculated using the leave-one-out cross-validation procedure. The final model used to predict the density of living trees across the study area included the number of detected treetops, slope inclination, SD of canopy gap area and the percentage of canopy gaps as explanatory variables (Table A1). The prediction produced by the final model had R^2^ = 0.38, RMSE = 129.94 and NMRSE = 0.17 (Fig. A2a). The histograms of the density of the trees observed and predicted in the study plots were visually similar, but without the last two histogram classes, i.e. the model underestimated the density of living trees in part of the study plot. (Fig. A2b). The living tree density using the final model was predicted on a raster grid with the calculated explanatory variables.

---

**Table A1.**
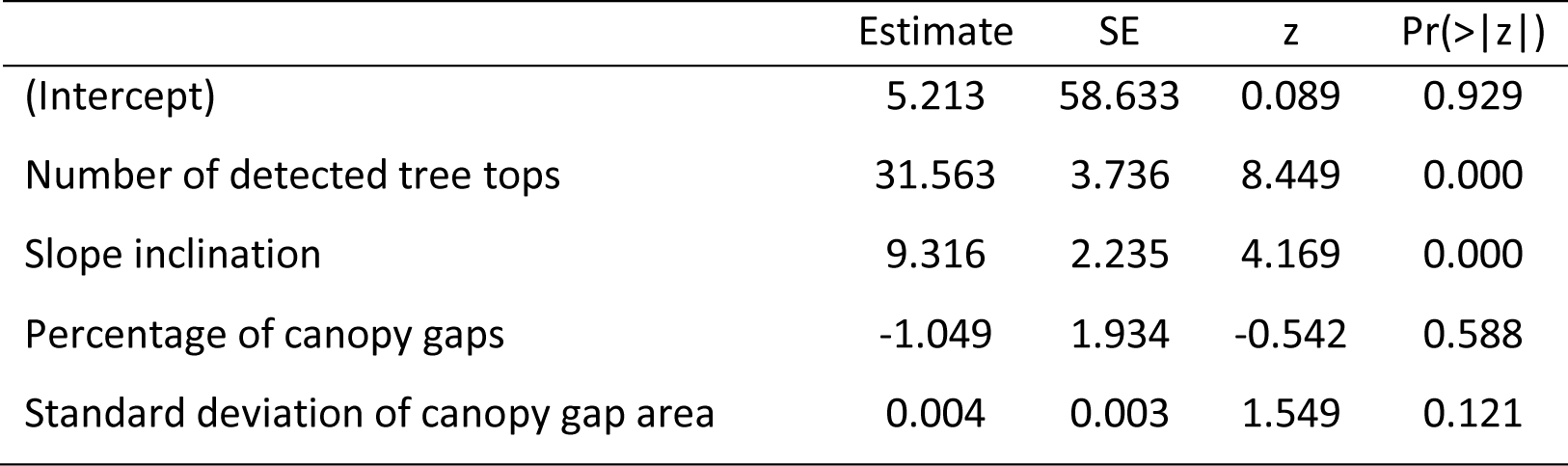
Final model describing the relationship between explanatory variables: number of detected treetops, slope inclination, percentage of canopy gaps and standard deviation of canopy gap area used to predict the density of living trees in the primeval European beech *Fagus sylvatica*-dominated forest in the Bieszczady Mountains (Poland) (N = 90).

**Fig. A1.**
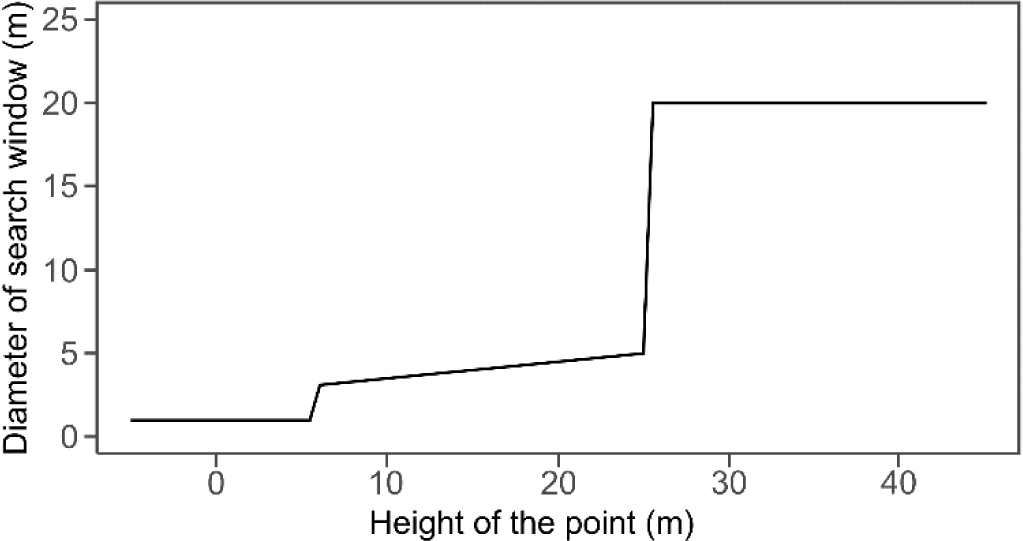
Relationship between the height of the detected treetops (m) and the search window diameter (m) used in the fixed-window-size local-maxima filtering method employed to detect treetops based on the Canopy Height Model acquired from Light Detection and Ranging point cloud data obtained as airborne laser scanning of the primeval European beech *Fagus sylvatica*-dominated forest in the Bieszczady Mountains (Poland).

**Fig. A2.**
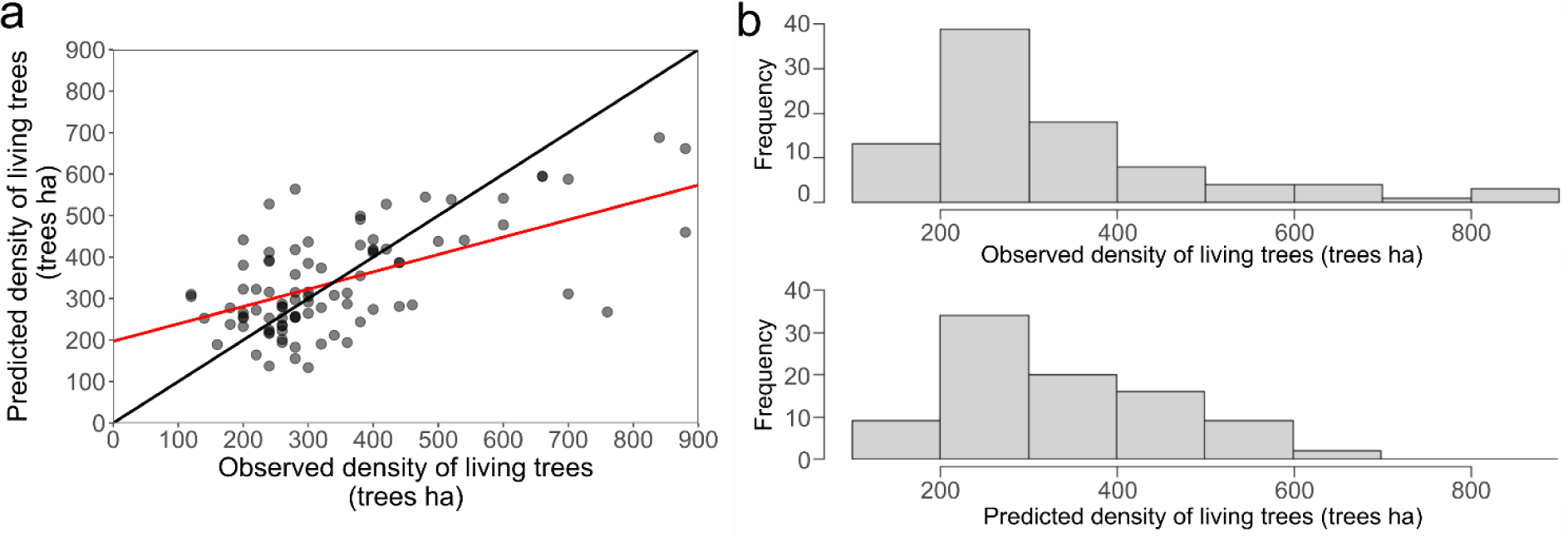
(a) Relationship between the predicted density of living trees (trees ha^-1^) and the observed density of living trees (trees ha^-1^) on the study plots in the primeval European beech *Fagus sylvatica*-dominated forest in the Bieszczady Mountains (Poland), and **(b)** histograms of the frequency distribution of the observed (upper row) and predicted (lower row) density of living trees. The black line in panel a indicates the theoretical linear relationship with a slope equal to 1 and the red line indicates the calculated linear relationship with a slope equal to 0.42 and an estimate equal to 197.21 trees ha^-1^.

